# Dynamic social envirotypes demonstrate adolescent brain dysconnectivity is contextual

**DOI:** 10.64898/2026.09.17.752338

**Authors:** Haily Merritt, Marlen Z. Gonzalez, Stephanie Noble, Andreia Sofia Teixeira

## Abstract

The adolescent social environment and dynamics therein have been linked to divergent brain organization and greater risk of psychopathology later in life, suggesting the neural underpinnings of mental health may depend on social context. To examine this, we leveraged longitudinal ABCD Study data to identify *dynamic social envirotypes*, or distinct trajectories of social environment change over four years. Dynamic social envirotypes differed in mental health outcomes, changes in mental health, and whole-brain functional connectivity, with each dynamic social envirotype exhibiting a characteristic pattern of brain network organization. Crucially, functional connectivity associations with internalizing and externalizing problems diverged across dynamic social envirotypes, with the highest and lowest social resource envirotypes showing largely opposing associations between connectivity and symptom severity. These findings suggest that *dysconnectivity is contextual* : there is no single brain connectivity pattern that is universally associated with better mental health. Rather, adolescent brain connectivity relates to mental health in context-specific ways, challenging normative narratives of the neural underpinnings of psychopathology and centering the role of context for understanding and treating mental health.

## Introduction

The social environment plays a key role in human health and wellbeing (Cohen, 1988, 2004; Krendl and Perry, 2021). Psychosocial adversity negatively impacts the function of many bodily systems (Vaidya et al., 2024), while social support buffers against the otherwise harmful influences of stress (Cohen and Wills, 1985). This influence of the social world on health and development is long-lasting, with early life adversity being linked to a number of psychopathologies later in life (Ho and King, 2021; Whittle et al., 2025), even when considering genetic influences (Caspi and Moffitt, 2006; Hicks et al., 2009). As an example, social support (or the lack thereof) in early childhood predicts internalizing and externalizing behaviors in kindergarten (Heberle et al., 2015). This pattern of results holds across focused samples (Rajaleid et al., 2016) and large data consortia (Dash et al., 2023), underscoring the importance of considering social context when investigating health and wellbeing. Critically, the social environment is dynamic and can be measured in numerous ways with overlapping and distinct psychopathological and neural associations (Rakesh et al., 2021d; Merritt et al., 2024), creating a research challenge which large consortia like the Adolescent Brain Cognitive Development Study (Garavan et al., 2018) are well-poised to address.

Adolescence in particular is a time of increased neural susceptibility to turbulence in the social environment (Foulkes and Blakemore, 2018; Michael et al., 2025; Branchi, 2011). This susceptibility motivates research to link psychopathology to its neurodevelopmental underpinnings, converging on several brain systems as key contributors to biomarkers of psychopathology (e.g. Default Mode Network (DMN), Dorsal Attention Network (DAN), and Limbic system) (Jirsaraie et al., 2025; Xie et al., 2023; Holz et al., 2023). Many results, however, have been directly contradictory, simultaneously proposing that, for example, increased and decreased DMN connectivity underlies depression (Prompiengchai and Dunlop, 2024). In the wake of ever more massive data consortia, the field is teeming with data to reconcile these contradictory findings, but it seems the volume of data alone is not the answer.

Advances in machine learning techniques have extracted high dimensional neural signatures of psychopathology (Xie et al., 2023; Drysdale et al., 2017), but these neural signatures do not always translate to clinical utility or relief for patients (Graham et al., 2019; Herrman et al., 2022). A potentially more effectual approach draws on developmental neuroscience (Tottenham, 2014), leveraging the interactions between context and neuroplasticity to elucidate the neuroscience of mental health (Merritt et al., 2026b; Michael et al., 2025). For example, in a cross-sectional study of 127 adolescents, Ramphal et al. (2020) reported that youth from more disadvantaged neighborhoods exhibited a positive association between fronto-amygdala connectivity and anxiety, whereas youth from less disadvantaged neighborhoods had a negative association. Similarly, Gee and colleagues found that amygdala-mPFC connectivity and anxiety symptoms had a positive association for maternally-deprived youth but a negative association for youth who did not experience maternal deprivation (Gee et al., 2013). Taken together, these studies suggest that *dysconnectivity is contextual* and, consequently, that effective treatment may depend on the environment. While network approaches have demonstrated the value of whole-brain perspectives on mental health (Lynch et al., 2024; Liu et al., 2025; Rakesh et al., 2021a,b, 2025b; Cocuzza et al., 2026), whether contextual dysconnectivity extends to other brain systems or across longitudinal data remains an open question. Addressing the contextuality of whole-brain dysconnectivity, while appreciating the high dimensionality and dynamicity of the social environment, therefore represents a critical challenge for human neuroscience and for the successful treatment of mental health.

To this end, we leveraged longitudinal data from 4,664 adolescents from the ABCD Study to characterize how social environments change over time. Our algorithm yielded three distinct trajectories of social environment experience, or *dynamic social envirotypes* (Merritt et al., 2026a): (1) a low social resource dynamic social envirotype, (2) a high social resource dynamic social envirotype, and (3) an intermediate social resource dynamic social envirotype. The high social resource dynamic social envirotype exhibited the best and most stable mental health, while the low social resource dynamic social envirotype experienced higher levels of psychopathology symptoms. The brain network connectivity also differed across these dynamic social envirotypes, with each exhibiting distinct patterns of connectivity. More consequently, we found that brain network connectivity that predicted better mental health diverged across the dynamic social envirotypes. More than that, the dynamic social envirotypes with the highest and lowest social resources showed largely opposing functional connectivity associations with internalizing and externalizing problems, demonstrating that dysconnectivity is contextual. Altogether, this work represents a critical contribution to the neuroscience of mental health, establishing developmental and ecological perspectives along with network science tools as necessary for investigating neural signatures of psychopathology.

## Results

To address the contextuality of whole-brain dysconnectivity while appreciating the high dimensionality and dynamicity of the social environment, we leverage the size and longitudinal nature of the Adolescent Brain Cognitive Development Study (Garavan et al., 2018). The first step is to identify the dominant modes of social environment change, which enables analysis of variability in mental health outcomes and in brain network organization across dynamic developmental contexts (see Figure 1). To do this, we extract clusters of trajectories of social environment experience, or *dynamic social envirotypes* (Merritt et al., 2026a). Given the critical role the adolescent social environment plays in the development of psychopathology, we assess outcomes of these dynamic social envirotypes on a suite of mental health measures. From a neuroecological and developmental perspective (Merritt et al., 2026b), we expect there to be differences in brain network organization across distinct social environment experiences due to calibration by the nervous system to environmental demands.

To test this, we contrast whole-brain connectivity across dynamic social envirotypes. Finally, we triangulate between brain, psychopathology, and social context to interrogate how dynamic social envirotypes contextualize the relationship between whole-brain connectivity and mental health.

### Dynamic social envirotypes capture modes of social environment change

How do social environments change over time? What are the dominant modes of social environment change (see Figure 1a)? To identify dynamic social envirotypes, we used 10 measures of the social environment recorded over four years: Parental Monitoring, Family Conflict (Youth-Report and Parent-Report), School Environment, School Involvement, School Disengagement, Independence, Religiosity, Prosociality, and Community Risk (see Figure 2a). These ten measures represent all indices of social environment quality that maximized response rate over time (*>* 60%). We opted to use more relational measures of the social environment as opposed to more structural measures like socioeconomic status (SES) since relational measures are more strongly linked to mental health (Coan et al., 2017; Danese and Widom, 2020) and because the relational measures have been much less investigated than SES (Merritt et al., 2026b; but see, for example, Rakesh et al., 2025b for relevant work with SES).

**Figure 1:**
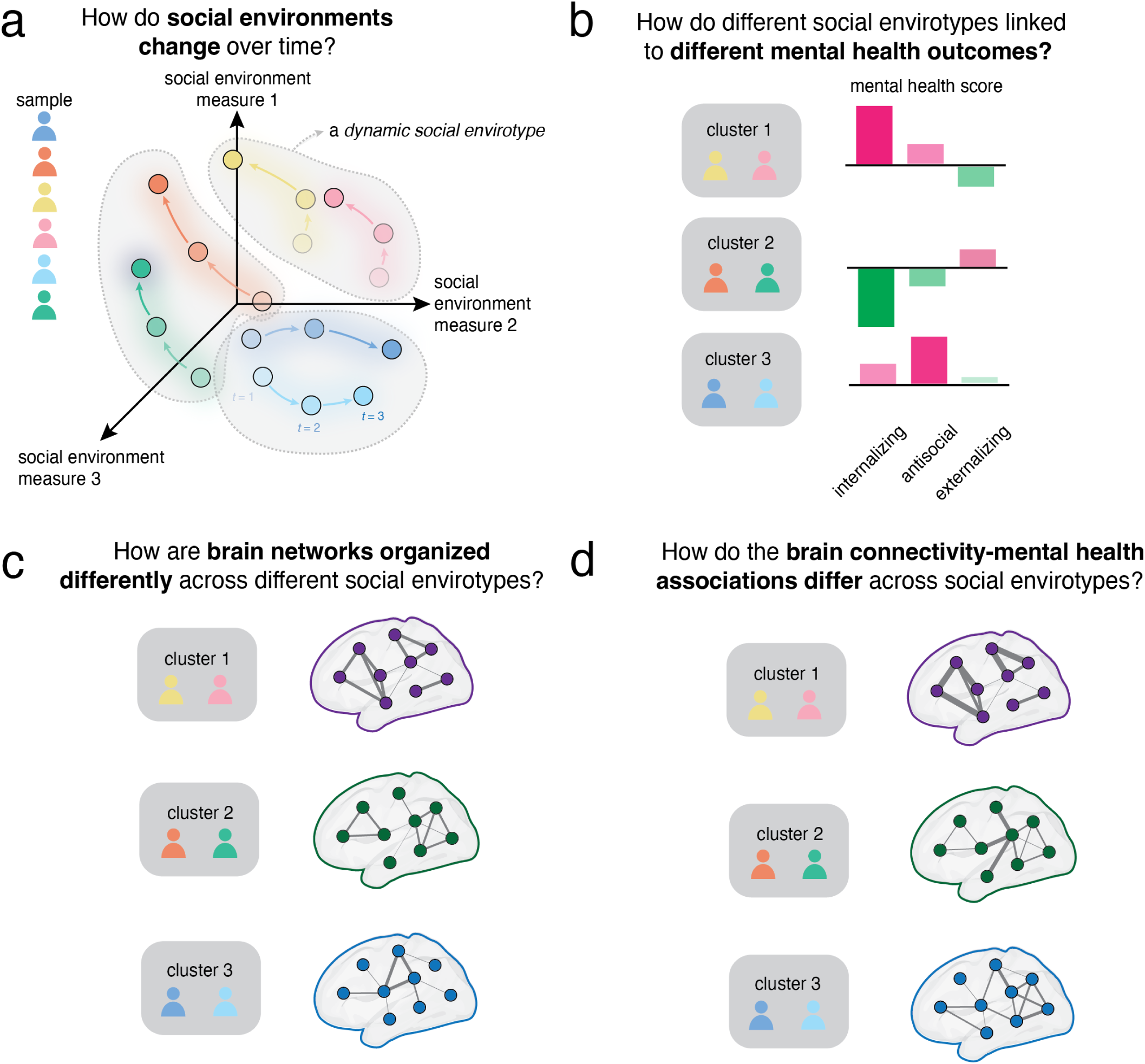
Schematic figure of analyses. We interrogate how social environments change over time using clustering analysis (panel a). For each cluster of trajectories of social environment change, which reflect underlying dynamic social envirotypes, we determine their mental health outcomes (panel b) and characteristic brain network organization (panel c). Finally, we assess contextual dysconnectivity, or how the brain network organization associated with better mental health looks different across different dynamic social envirotypes (panel d).

**Figure 2:**
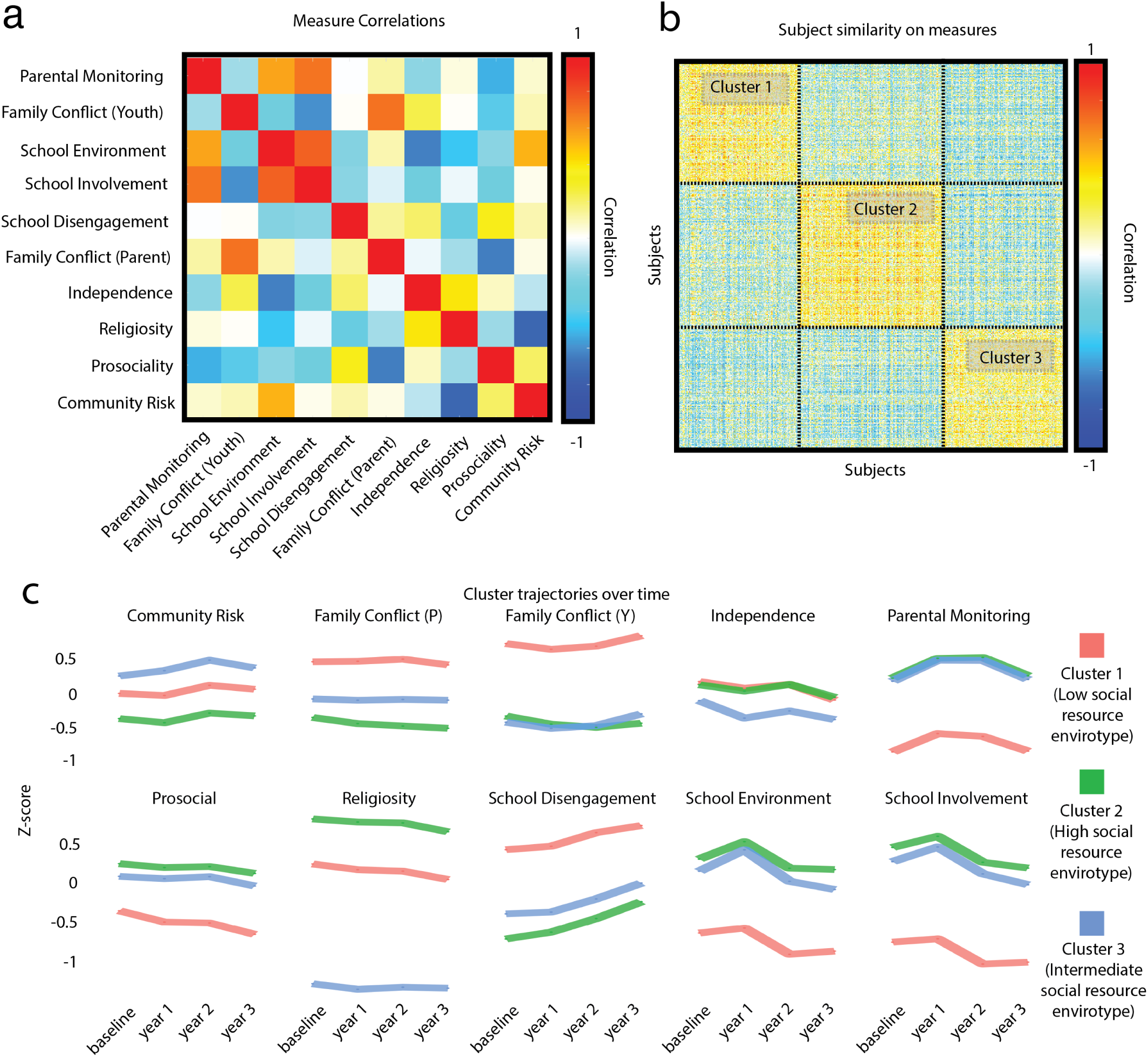
Measures of the Social Environment & Cluster Trajectories. We identified the dynamic social envirotypes using a suite of 10 longitudinal measures of the social environment. Panel (a) shows the inter-measure similarity across subjects. Each subject is represented by a 40 × 1 vector (10 measures at 4 time points), so to calculate subject similarity we compute the correlations between all pairs of subjects using these vectors. Panel (b) displays the resulting similarity matrix, with the rows and columns ordered to indicate the three primary envirotypes that result from our clustering algorithm. The trajectories of each resulting envirotype on each measure are shown in panel (c).

Using these ten measures collected across four time points, we clustered the *N* = 4664 adolescents who had complete data. Our clustering algorithm yielded five clusters: *N*_1_ = 1465*, N*_2_ = 1741*, N*_3_ = 1449*, N*_4_ = 5, and *N*_5_ = 4 (see Figure 2b). We did not select the number of clusters *a priori*; rather, our 1000 iterations of clustering and validation analysis showed that this cluster composition reflects the most stable clusters and is sufficiently representative of the dominant modes of social environment change in the data (see Supplementary Materials and Supplementary Figure S1 for details). In order for our subsequent analyses to be sufficiently statistically powered to detect meaningful differences, we focus our statistical analyses on Clusters 1, 2, and 3, which represent 99.8% of the sample used for clustering. These three clusters exhibit statistically distinguishable dynamic social envirotypes (see Figure 2c, Supplementary Table S2 for all *p*-values of cluster-wise comparisons, Supplementary Table S3 for all *p*-values of temporal comparisons, and Supplementary Table S4 for effect sizes). Cluster 3, which exhibited a dynamic social envirotype characterized by intermediate social resources, changed the most over time, notably between baseline and year 1 and between years 2 and 3. Cluster 1, which had a relatively low social resources, and Cluster 2, which had the most resourced dynamic social envirotype, were both more stable but displayed more change across social environment measures between years 2 and 3. Cluster 1 differed significantly from both Clusters 2 and 3 on all measures at all time points, except for Independence where it was indistinguishable from Cluster 2 (all *p*s *<* 0.00001). Clusters 2 and 3 differed significantly on most measures at most time points, with Cluster 2 more consistently exhibiting a dynamic social envirotype with more social resources. Hereafter, as a shorthand we refer to Cluster 1 as the low social resource envirotype, Cluster 2 as the high social resource envirotype, and Cluster 3 as the intermediate social resource envirotype.

We also assessed potential confounding variables on the dynamic social envirotypes, including parental income, parental education, pubertal status, race/ethnicity, and age. We found that the intermediate social resource envirotype had higher income and education than the high and low social resource envirotypes (*p*s *<* 0.00001), lower pubertal status than the high and low social resource envirotypes(*p* = 0.001), and a lower proportion of Black or African American (*p* = 1.7 × 10*^−^*^12^) and Hispanic or Latino (*p* = 2.3 × 10*^−^*^8^) youths. There was no difference in age between any of the envirotypes, nor were there any statistically significant differences between the high and low social resource envirotypes on any of these measures (see Supplementary Figure S3). Additionally, the timing of data collection with respect to COVID-19 lockdowns was not distributed differently across envirotypes (*χ*^2^(2) = 4.05*, p* = 0.132; see Supplementary Figure S4).

Generally, the low social resource envirotype was characterized by higher conflict and disengagement, with more change taking place in later years. The high social resource envirotype exhibited lower risk and lower conflict, but also saw more change in later years. The intermediate social resource envirotype had the most change and experienced lower religiosity, higher risk, lower independence, and a higher socioeconomic status.

### Dynamic social envirotypes distinguish adolescent mental health outcomes and changes

To what extent are these dynamic social envirotypes associated with distinct mental health outcomes and changes (see Figure 1b)? To address this, we examined differences in mental health at baseline, outcomes (Year 2), and changes. Looking at mental health scores at baseline, we found that all pairwise comparisons for all measures were significant even when controlling for SES (except Personal Strength, on which high and intermediate social resource envirotypes did not differ; all other *p*s < 0.00001; see Supplementary Table S5 for all *p* values and Supplementary Table S6 for all effect sizes). The low social resource envirotype had the worst mental health scores, while the high social resource envirotype had the best mental health scores (see Supplementary Figure S5). At Year 2 (i.e., the third time point), all pairwise comparisons for all measures were significant (all *p*s *<* 0.002), such that the low social resource envirotype again had worse mental health outcomes while the high social resource envirotype had better mental health outcomes (see Figure 3; see Supplementary Table S5 for all *p* values and Supplementary Table S6 for effect sizes).

**Figure 3:**
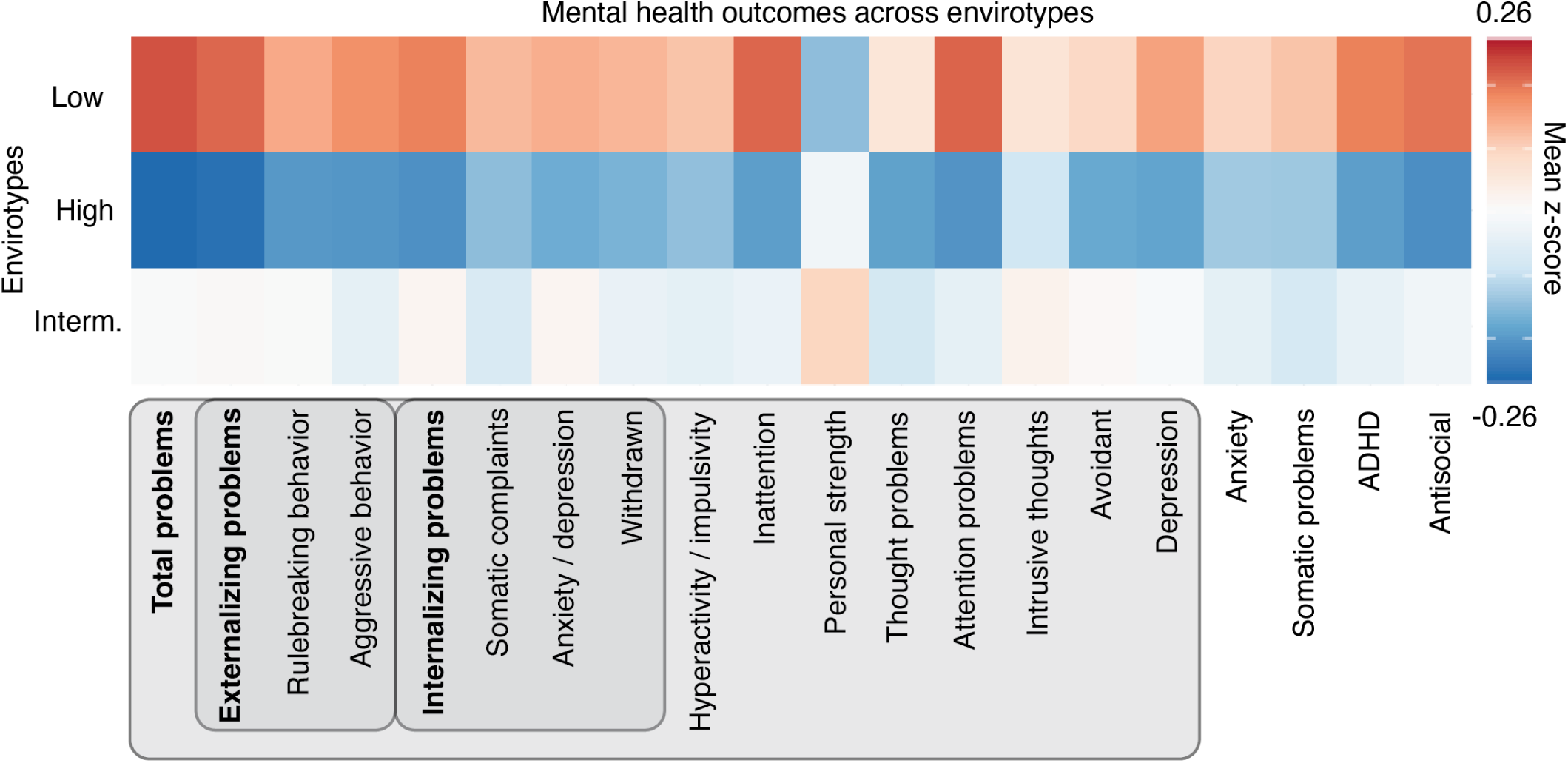
Dynamic social envirotypes distinguish mental health outcomes. Measures (taken during Year 2) used to compute composites are indicated by the boxes, with the composite measure’s name in bold. All measures are z-scored to be on the same scale. Higher scores indicate greater endorsement of symptoms, so worse mental health (except for Personal Strength). For each mental health measure, all pairwise comparisons between envirotypes are significantly different (all *p*s *<* 0.00001).

When looking at changes in mental health between clusters, we found that all pairwise comparisons were significantly different for all measures (all *p*s *≤* 0.0001; see Supplementary Figure S6; see Supplementary Table S5 for all *p* values and Supplementary Table S6 for effect sizes). More specifically, the low social resource envirotype exhibited the biggest improvement in symptoms of ADHD, Aggressive Behavior, Anxiety, Hyperactivity/Impulsivity, and Thought Problems and the biggest decline in symptoms of Attention Problems, Inattention, Somatic Complaints, and Somatic Problems; the high social resource envirotype showed the biggest improvement in symptoms of Externalizing Problems, Personal Strength, and Rulebreaking Behavior and the biggest decline in symptoms of Withdrawn Behavior; and the intermediate social resource envirotype had the biggest improvement in symptoms of Avoidant Behavior and the biggest declines in symptoms of ADHD, Antisocial Behavior, Anxiety/Depression, Anxiety, Total Problems, Depression, Externalizing Problems, Internalizing Problems, and Rulebreaking Behavior.

Additionally, we leveraged the fact that social environment measures were collected annually while mental health measures were collected biannually to assess which changes in the social environment were associated with changes in mental health measures. After correcting for multiple comparisons, there were no social environment changes that were associated with significantly improved or worsened mental health outcomes (see Supplementary Figure S7 for uncorrected results).

In sum, we found that the low social resource envirotype experienced more symptoms of psychopathology across a range of measures. The best mental health outcomes were associated with the high social resource envirotype.

### Dynamic social envirotypes exhibit characteristic brain network organization

How are brain networks organized differently across the dynamic social envirotypes (see Figure 1c)? We identified characteristic connectivity for each cluster, with differences distributed across the brain for all pairwise envirotype comparisons when controlling for SES (see Figure 4). In particular, the low social resource envirotype exhibited higher connectivity between Vis and association systems (e.g. FP, RT, VAN; see Supplementary Table S7 for longer names of brain system abbreviations), in addition to lower connectivity between Aud and each of VAN and CO and between SMh and Sal. This envirotype also exhibited higher connectivity within CP and lower connectivity within CO. The high social resource envirotype, on the other hand, had higher connectivity between Aud and CP, between SMh and both RT and Sal, between SMm and each of Sal, VAN, and CP, within Vis and FP, as well as lower connectivity between Vis and each of VAN, DMN, and Aud and between SMm and DAN. The intermediate social resource envirotype had higher connectivity within CO and DMN but lower connectivity between these systems, between DMN and DAN, and between CO and each of CP, FP, and Sal. Additionally, this envirotype had higher connectivity between SMh and VAN but lower connectivity between SMh and CP, between SMm and each of FP and Aud, and within VAN and RT (see Figure 4 for envirotypical FC according to test statistics, Supplementary Figure S8 for the node- and system-level mean FC for each envirotype, Supplementary Figure S9 for the test statistics, and Supplementary Figure S10 for effect sizes of these tests). In general, differences in connectivity across envirotypes were distributed across the brain and hint at potential trade-offs in the connectivity between sensorimotor and association systems across envelopes.

**Figure 4:**
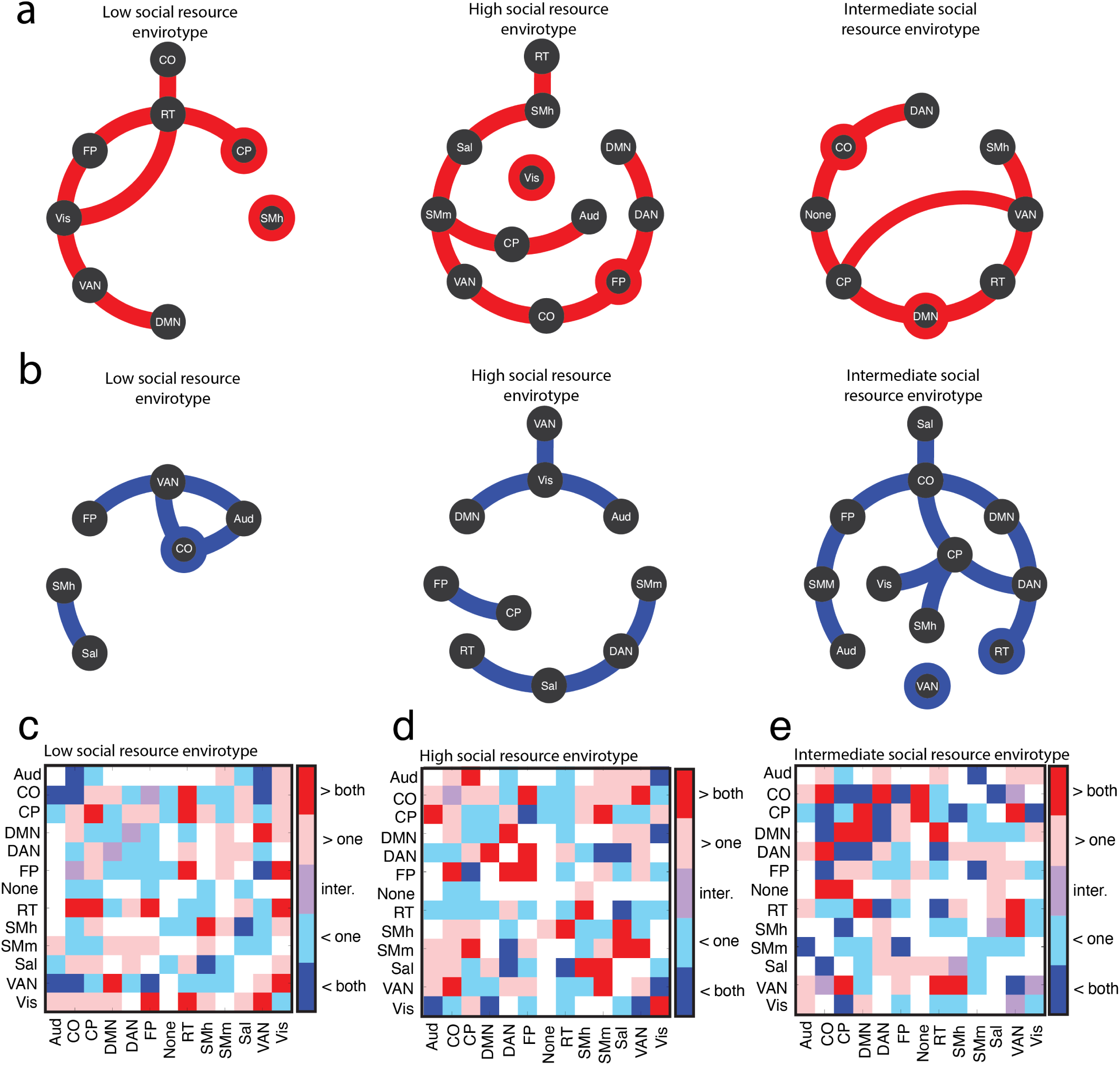
Dynamic social envirotypes exhibit characteristic functional connectivity. Panels (a-c) (and (d-f)) show the system-wise connections for each envirotype that were significantly higher (and lower) than the other two envirotypes. Brain systems are ordered differently in each envirotype’s plot for visual clarity, but panels (g-i) show all system-wise comparisons of connectivity with identical ordering for all envirotypes. In our categorical colorbar, bright red (blue) indicates higher (lower) connectivity than both other envirotypes and is highlighted in panels (a-f). Light red (blue) indicates higher (lower) connectivity than just one other envirotype, purple indicates statistically intermediate connectivity (i.e., statistically lower connectivity than one envirotype but statistically higher connectivity than the other), ^1^a^1^nd white represents no significant differences.

### Dynamic social envirotypes emphasize the role of context in associations between brain connectivity and mental health

How do the brain connectivity-mental health associations differ across social envirotypes (see Figure 1d)? We also investigated how the brain network organization associated with better mental health–focusing on symptoms of Internalizing and Externalizing Problems–looks different for the distinct dynamic social envirotypes, controlling for SES (see Supplementary Figure S11 for this association for two exemplary edge weights). We identified characteristic associations between brain connectivity and mental health for each cluster, with nearly opposite patterns with respect to the low and high social resource envirotypes (see Figure 5). For symptoms of Internalizing Problems, the low social resource envirotype’s strongest associations included higher connectivity within DMN, Vis, None, DAN, and CP and lower connectivity within VAN, Sal, and RT. In terms of connectivity between systems, this envirotype had lower connectivity between VAN and Vis, SMh and CP, DAN and CP, RT and CP, None and Sal, and RT and Sal linked to increased symptoms of Internalizing Problems. The only higher between system connectivity was between DMN and Sal, an association with Internalizing Problems shared with the intermediate social resource envirotype but reversed in the high social resource envirotype. The strong associations with symptoms of Internalizing Problems for the high social resource envirotype also had lower connectivity between Sal and CO, Sal and Vis, and Aud and SMm but higher connectivity between RT and DAN and between CP and Aud. Associations for the intermediate social resource envirotype exhibited higher connectivity between DMN and Sal, RT and CP, and VAN and DAN but lower connectivity between Sal and FP, FP and RT, CP and SMh, and SMh and DAN.

**Figure 5:**
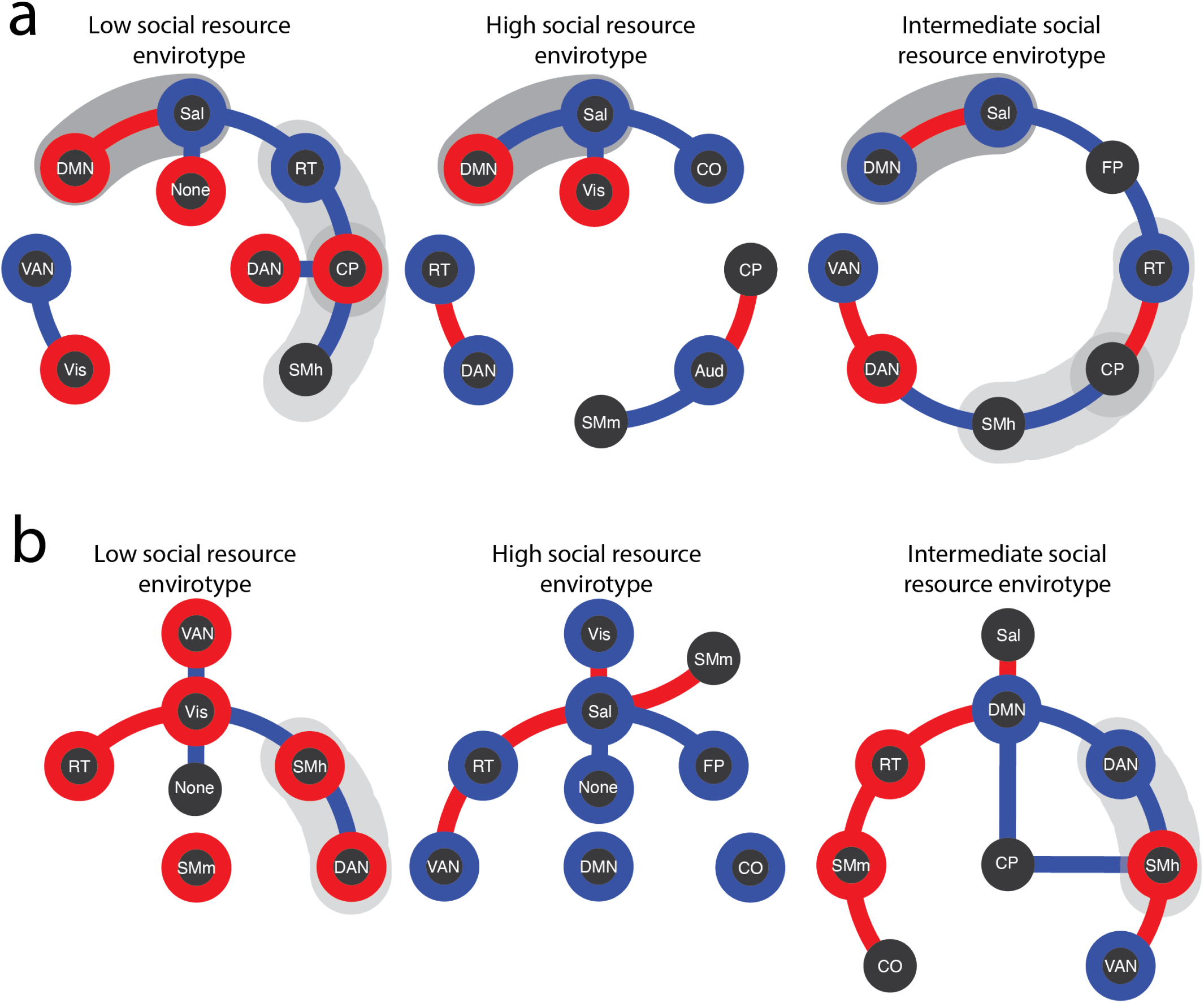
Healthy brain network organization varies across dynamic social envirotypes. We assess the extent to which internalizing problems (row a) and externalizing problems (row b) are associated with distinguishable patterns of connectivity for the different envirotypes. In both cases, red indicates a significant positive association (higher connectivity is associated with more symptoms), while blue indicates a significant negative association (higher connectivity is associated with fewer symptoms). Red (blue) outlines around nodes represent significant positive (negative) associations between within system connectivity and symptoms. Gray outlines around connections indicate shared neural associations across envirotypes. The connections shown here represent the 10% strongest associations for each cluster for each measure. Additional associations can be found in Supplementary Figure S13 and Supplementary Figure S14.

For Externalizing symptoms, we observed a stark dissociation in the links to within-system connectivity across envirotypes. For the high social resource envirotype, we saw lower within-system connectivity was associated with more symptoms of Externalizing Problems, while for the low social resource envirotype the association was reversed. Put quantitatively, all nine systems that had significant associations between their within-system connectivity and externalizing problems for both the low and high social resource envirotypes flipped the sign of their association between envirotypes (i.e., Aud, CO, DMN, DAN, FP, RT, Sal, VAN, Vis). The intermediate social resource envirotype, on the other hand, exhibited less of a clear-cut association. For both the low and intermediate social resource envirotypes, higher connectivity within somatomotor systems was linked to more externalizing symptoms, while the high social resource envirotype showed the inverse association (see Supplementary Figure S12 for edge-level *t* statistics, Supplementary Figure S13 for systemwise *Z* statistics, Supplementary Figure S14 for system-level summaries of envirotypical associations, Supplementary Figure S15 for effect sizes of tests predicting Externalizing Problems, and Supplementary Figure S16 for effect sizes of tests predicting Internalizing Problems). In nearly all of these cases (except the positive association between intra-SMh connectivity and symptoms), the neural connectivity patterns associated with increased psychopathology diverge from the group pattern norm for that dynamic social envirotype. Overall, these findings suggest that *dysconnectivity is contextual*.

## Discussion

We leveraged longitudinal social environment data from thousands of adolescents to identify three distinct dynamic social envirotypes, which reflect the dominant modes of social environment change. The high social resource envirotype exhibited the least symptomatic and most stable mental health, while the low social resource envirotype experienced the most symptoms of psychopathology. Each dynamic social envirotype exhibited characteristic brain network connectivity in such a way that suggests a potential trade-off in connectivity between and within somatosensory and association areas. Finally, our approach revealed evidence for contextual dysconnectivity: the brain network connectivity that was associated with better mental health looked different across the dynamic social envirotypes, emphasizing the importance of considering context when studying the neural underpinnings of mental health.

Despite using different methods than previous social and developmental neuroscience research, our findings are largely consistent with the broader body of literature linking the social environment to mental health. More specifically, like Heberle et al. (2015); Rajaleid et al. (2016); Dash et al. (2023); Cohen (2004); Antonucci et al. (2014), we find that more socially resourced environments tend to be linked to better mental health outcomes, while more adverse environments are linked to worse outcomes. Although theoretical work emphasizes the distinction between different dimensions of the environment–such as harshness/threat, deprivation, or instability/unpredictability (Sheridan and McLaughlin, 2014; McLaughlin et al., 2014)–, data-driven approaches tell a subtly different story. For example, Merritt and colleagues examine longitudinal clusters using a similar approach, identifying four clusters that differ based on the quality and stability of the social environment (Merritt et al., 2026a). Their clusters seem to exhibit trade-offs in mental health outcomes, while ours are more clearly stratified. This could be due to the different measures used to cluster, or it could signal the importance of social environment quality–again clearly stratified in our clusters–over other dimensions for mental health.

Indeed, many studies have leveraged the size of the ABCD study to apply clustering techniques. Wang et al. (2023), Lichenstein et al. (2022), and DeRosa et al. (2024) cluster the neuroimaging data itself to identify neuroendophenotypes, which they then link to psychopathology, cognition, and sociodemographic features. Both Wang et al. (2023) and DeRosa et al. (2024) identify subgroups that exhibit both higher levels of psychopathology and increased within-system functional connectivity, similar to our Cluster 1 (low social resource envirotype). Fekson et al. (2023), on the other hand, performs latent profile analysis on inhibitory control measures, then links these profiles to intra-DMN connectivity. Together, these studies emphasize that there are indeed distinct patterns of brain organization. Our approach moves beyond analyzing associations of subtypes to consider the *implications of these distinctions*. We contextualize not just differences in functional connectivity itself but the functional connectivity linked to better mental health across different social environments. Moreover, by clustering individuals based on longitudinal social environment measures, our work builds on the rich history of research in developmental and ecological neuroscience, enabling a clearer mechanistic understanding of the neuroscience of mental health (Merritt et al., 2026b).

One of the most common indices of social environment quality is socioeconomic status (SES), which is often operationalized as (parent) income or education and has been linked to better mental health (Rakesh and Whittle, 2021; Webb et al., 2022). Notably, the dynamic social envirotype with the highest parent income and education (intermediate social resource envirotype) does not exhibit the best mental health outcomes. While the high social resource envirotype has a lower SES, they exhibit higher religiosity than the other two envirotypes, which previous work has shown is tightly linked to better mental health (Wong et al., 2006). Additionally, this higher SES cluster exhibits higher Community Risk, which in this dataset indexes the accessibility of alcohol and other substances, and lower Independence. Given the link between substance use and psychopathology (Voepel-Lewis et al., 2025) and between independence or autonomy and wellbeing (Ruiz and Yabut, 2024), it may be that any benefits of SES to mental health are dampened by these other social factors. This would echo work emphasizing that various facets of the social environment interact in multitudinous ways (Danese and Widom, 2020; Webb et al., 2022; Rakesh et al., 2021d; Antonucci et al., 2014), making clarity of measurement critical for research moving forward.

In our brain network analyses, we found that the patterns of brain connectivity that were characteristic for each dynamic social envirotype were dissociable, meaning brain network organization looks different given different experiences, echoing previous subtyping approaches as described above. Importantly, the low and high social resource envirotypes have opposite mental health outcomes, but this does not necessarily entail that their patterns of dysconnectivity will be inverted (Merritt et al., 2026b). More than that, this result is clearly in line with ecological perspectives on neurodevelopment (Merritt et al., 2026b). That is, while connectivity differences between envirotypes were distributed across the brain, thereby emphasizing the importance of whole-brain and network approaches, a subset of systems emerged as especially important and pointed toward potential trade-offs in connectivity depending on the environment. In particular, the intermediate social resource envirotype exhibited higher connectivity within CO and DMN, but the low and high social resource envirotypes had higher connectivity between these systems. A number of studies have identified higher intra-DMN connectivity in more socially supportive environments (Mwilambwe-Tshilobo et al., 2019, 2022; Spreng et al., 2020; Merritt et al., 2026a).

Relatedly, the low social resource envirotype had higher connectivity between Vis and many association systems, while the high social resource envirotype had higher connectivity within Vis. Similarly, the low social resource envirotype had higher connectivity within SMh but the high and intermediate social resource envirotypes displayed higher connectivity between this system and association systems. While differences in somatosensory connectivity have been reported by a number of studies (Rakesh et al., 2021c; Merritt et al., 2026a; Mwilambwe-Tshilobo et al., 2019), theoretical perspectives have yet to make sense of this finding. A recent article proposes two putative pathways, one more protracted route via stress physiology and one more acute via supportive touch (Merritt et al., 2026b), but the precise mechanism remains undetermined. Clarifying the mechanisms underlying connectivity-based calibrations to the environment may also elucidate the extent to which such calibrations require yoked trade-offs in connectivity within and between systems.

Notably, we found evidence for contextual dysconnectivity. Specifically, for the dynamic social envirotype with fewer social resources, higher within-system functional connectivity was linked to more externalizing problems, whereas the higher social resourced dynamic social envirotype exhibited the reverse association. These findings add to a body of work presenting a resounding challenge to research seeking a unified or universal neural biomarker for psychopathology (Xie et al., 2023). Previous research has revealed divergent neural underpinnings of anxiety (Ramphal et al., 2020; Gee et al., 2013), substance abuse (Rakesh et al., 2021a), conduct problems (Liu et al., 2025), and mental health problems in general (Rakesh et al., 2025b) across groups with different social experiences (e.g. maternal deprivation or socioeconomic status). More than there being neural variability associated with psychopathology (Segal et al., 2025), dysconnectivity is contextual. Ecological and developmental perspectives offer an explanation for this.

The plastic developing brain adapts to the environment it is in–whether adverse or enriching–, modifying its structure, organization, and function to suit the pressures and demands of its context in accordance with genetic constraints (Tottenham, 2014; Sheridan and McLaughlin, 2014; McLaughlin et al., 2014; Ellis and Del Giudice, 2014; Michael et al., 2025). A consequence of these processes of plasticity and adaptation is that across different environments, there are distinguishable patterns of brain network functional connectivity (Merritt et al., 2026b,a). Importantly, there are many routes of brain network reconfiguration (Rakesh et al., 2021a; Vanes et al., 2025; Rakesh et al., 2025a). Despite universal material constraints (Bullmore and Sporns, 2012), brain networks may exhibit multifinality in their organization, so it is possible that two relatively dissimilar environments could yield relatively similar network neuroendophenotypes. For this reason, research simply investigating whether there are brain connectivity differences between different social environments or between subtypes is telling only part of the story (Merritt et al., 2026b). Necessary to the narrative is an understanding of ‘fitness’ or ‘adaptedness’ to the environment, especially given that recent work shows that environmental mismatch increases “neuroburden’ (Lee and Gonzalez, 2025). Mental health outcomes are one way to operationalize ‘fitness’ or ‘adaptedness.’ Moving forward, research on the neuroscience of mental health will benefit from a triangulation approach, situating brain organization and measures of psychopathology in their (social) context.

Despite the advances to our understanding of the neuroscience of mental health, this work has three key limitations that call for future work. First, activation and seed-based connectivity studies have made clear that limbic connectivity, such as the amygdala-prefrontal cortex circuit, plays a central role in the neuroscience of mental health (Tottenham, 2014; Gee et al., 2013; Ramphal et al., 2020; Morawetz et al., 2021). When using whole-brain approaches, the signal quality from midline nuclei and limbic structures unfortunately decreases, limiting the extent to which whole-brain research, despite its advantages, can be thoroughly integrated with other theory-driven and animal model-inspired work. To clarify mechanistic interpretations, dialogue between whole-brain and focal studies is crucial. Second, while our holistic clustering approach identifies data-driven differences in social environment experience, several studies have demonstrated there is much variability in social measures’ associations with connectivity, including for operationalizations of the same construct (Rakesh et al., 2021d; Merritt et al., 2024). Our approach is unable to capture such nuance. For this reason, there is value in combining data-driven clustering studies with approaches focusing on the nuances of particular measures of interest. Finally, with the increasing availability of longitudinal population neuroimaging datasets like ABCD, researchers have been able to more effectively address questions regarding variability in the pace, organization, and trajectory of neurodevelopment (Tooley et al., 2021; Rakesh et al., 2023; Sydnor et al., 2023). Emerging work suggests there may not be a single trajectory along which functional connectivity develops (Rakesh et al., 2021a, 2025a, 2023). Instead, the plasticity and adaptation that are central to neurodevelopment (Michael et al., 2025) may entail divergent trajectories of neurodevelopment in accordance with experience (Merritt et al., 2026b). Critically, this would challenge our understanding of normative neurodevelopment and its psychopathological consequences. Therefore, characterizing and contextualizing such trajectories will be an especially fruitful endeavor for future research.

## Conclusions

We identified distinct dynamic social envirotypes with divergent mental health outcomes. The brain organization was dissociable across these dynamic social envirotypes, but, more impactfully, the brain organization associated with better mental health looked different across the different dynamic social envirotypes. This result underscores that dysconnectivity is contextual and emphasizes the importance of a triangulation approach for the neuroscience of mental health. Moving forward, techniques that can elucidate the mechanisms for contextualized connectivity calibration and that can characterize divergent trajectories of neurodevelopment across contexts will be extremely valuable.

## Code and Data Availability

All data were obtained from the Adolescent Brain Cognitive Development Study (see below). Code used for analyses can be found at https://github.com/h-merritt/dynamic_social_envirotypes.

## Acknowledgements

Data used in the preparation of this article were obtained from the Adolescent Brain Cognitive DevelopmentSM (ABCD) Study (https://abcdstudy.org), held in the NIMH Data Archive (NDA). This is a multisite, longitudinal study designed to recruit more than 10,000 children age 9-10 and follow them over 10 years into early adulthood. The ABCD data repository grows and changes over time. The ABCD data used in this report came from NBDC Digital Object Identifier (DOI). DOIs can be found at https://www.nbdc-datahub.org/abcd-study. The ABCD Study® is supported by the National Institutes of Health and additional federal partners under award numbers U01DA041048, U01DA050989, U01DA051016, U01DA041022, U01DA051018, U01DA051037, U01DA050987, U01DA041174, U01DA041106, U01DA041117, U01DA041028, U01DA041134, U01DA050988, U01DA051039, U01DA041156, U01DA041025, U01DA041120, U01DA051038, U01DA041148, U01DA041093, U01DA041089, U24DA041123, U24DA041147. A full list of supporters is available at https://abcdstudy.org/federal-partners.html. A listing of participating sites and a complete listing of the study investigators can be found at https://abcdstudy.org/consortium_members/. ABCD consortium investigators designed and implemented the study and/or provided data but did not necessarily participate in the analysis or writing of this report. This manuscript reflects the views of the authors and may not reflect the opinions or views of the NIH or ABCD consortium investigators.

## Competing interests

The authors have no competing interests to declare.

## Methods

### Data

We used ABCD BIDS version 1.2.0 minimally pre-processed by the DCAN lab (additional information about the data pre-processing can be found in the Supplemental Materials). We note that this data release includes only one timepoint of neuroimaging data. From this release, we used social environment quality data over four years and resting state fMRI data at baseline from 4664 (47% female) adolescents age 9.98 *±* 0.62 years in the Adolescent Brain Cognitive Development (ABCD) dataset Garavan et al. (2018) (see Supplementary Figure S2 for the sample size with each inclusion step). This study was approved by the Institutional Review Board at each study site, with centralized IRB approval from the University of California, San Diego. Informed consent and assent were obtained from all parents and children, respectively. Our social envirotyping data included ten youth-report social environment quality measures: Parental Monitoring, Family Conflict (Youth-Report and Parent-Report), School Environment, School Involvement, School Disengagement, Independence, Religiosity, Prosociality, and Community Risk. We used these specific measures because they represented all of the measures of social environment quality with at least a 60% response rate across four time points. While a higher response rate could be achieved by focusing on fewer time points (e.g. 80% across three time points), we were especially interested in change over time and therefore opted to maximize the number of time points. We note that while objective measures like socioeconomic status have been widely used to examine brain-environment associations (e.g. Rakesh et al. (2021d); Ellwood-Lowe et al. (2021)), recent work has suggested that subjective indicators of social environment quality may have greater bearing on mental health and wellbeing Coan et al. (2017); Danese and Widom (2020); Amieva et al. (2010). All measures were z-scored to be on the same scale. See Supplementary Table S1 for sample demographic information.

Details of MRI acquisition and pre-processing for ABCD data have been described elsewhere Casey et al. (2018); we include details relevant to this study in the Supplementary Materials. We note that scans were typically completed on the same day as the social environment surveys, but could also be completed at a second testing session. To construct the functional connectivity matrices using the parcellated time series for 333 cortical regions, we computed the Pearson correlation between the time series of all pairs of nodes for each subject. This procedure yielded a 333 × 333 functional connectivity matrix for each subject.

### Identifying dynamic social envirotypes using clustering

To identify dynamic social envirotypes, we applied a clustering algorithm to the social environment data. Specifically, we first concatenated the scores for each subject, so that we had a 40 × 1 vector (10 measures at four time points) for each subject. We computed the pairwise correlation matrix between all pairs of subjects using these vectors. We applied an arctangent transformation to this matrix so all values were greater than 0, which our clustering algorithm requires. To partition this similarity matrix (and thereby participants) into clusters, we used the Louvain algorithm for modularity maximization, as in (Merritt et al., 2026a). A description of the algorithm is provided in the Supplementary Materials. Briefly, this procedure yields clusters of participants whose similarity to each other maximally exceeds what would be expected by chance. The advantage of this procedure is that it is fully data-driven; it requires no *a priori* specification of the number of clusters, the size of clusters, or any patterns of change or dimensions of the measures that may be of interest. We validated that our identified clusters were meaningfully representative of their dynamic social envirotypes. This procedure is described in detail in the Supplementary Materials. Following validation, we perform ANOVAs and *post hoc* tests where appropriate to determine differences between clusters and over time across the social environment measures, correcting for multiple comparisons using the Benjamini-Hochberg technique. Additionally, we considered whether the potential confounds of parental income, parental education, pubertal status, age, or race/ethnicity could account for cluster composition. Finally, given the timing of data collection for this release of ABCD data, we assessed whether the timing of COVID-19 lockdowns was differently distributed across clusters using a *χ*^2^-test, since COVID is known to influence social support and mental health (Xiao et al., 2021).

### Assessing mental health outcomes across different dynamic social envirotypes

We determined the extent to which the dynamic social envirotypes exhibited divergent mental health outcomes using a suite of 17 mental health measures: Somatic Complaints, Rulebreaking behavior, Aggressive behavior, Anxiety / Depression, Withdrawn Depression, Hyperactivity / Impulsivity, Inattention, Personal strength, Thought problems, Attention problems, Intrusive thoughts, Avoidant behavior, Depression, Anxiety (DSM), Somatic problems (DSM), ADHD (DSM), and Antisocial behavior (DSM). Additionally, we included three composite measures: Externalizing problems (which includes Rulebreaking behavior and Aggressive behavior), Internalizing problems (which includes Somatic complaints, Anxiety / Depression, and Withdrawn depression) and Total problems (which includes Somatic complaints, Rulebreaking behavior, Aggressive behavior, Anxiety / Depression, Withdrawn depression, Hyperactivity / Impulsivity, Inattention, Personal Strength, Thought problems, Attention problems, Intrusive thoughts, Avoidant behavior, and Depression). We assessed mental health differences in three ways: (1) differences in mental health scores between envirotypes at baseline, (2) differences in mental health scores between envirotypes at Year 2, and (3) differences between envirotypes in the amount of change in mental health scores between baseline and Year 2. In all three cases, we performed ANOVAs and *post hoc* tests where appropriate, controlled for SES given its close relationship to our social measures, and corrected for multiple comparisons.

Social environment measures were collected annually while mental health measures were collected biannually. To leverage this schedule, we assessed whether changes in social environment measures were associated with concurrent changes in mental health for each dynamic social envirotype. To do this, we first calculated each individual’s difference in differences score using the social environment measures (*t*_3_ *− t*_2_) *−* (*t*_2_ *− t*_1_), comparing across envirotypes. We used these scores to predict each individual’s mental health score at *t*_3_ using linear regression, comparing across envirotypes and correcting for multiple comparisons using the Benjamini-Hochberg technique.

### Examining differences in brain network organization between dynamic social envirotypes

We subsequently compared dynamic social envirotypes based on their whole-brain functional connectivity (FC; N = 333 cortical regions of interest using the Gordon parcellation Gordon et al. (2016)). First, we performed mass *t* -tests on the edge weights (i.e., the correlation coefficients of the time series of the nodes), correcting for multiple comparisons using the Benjamini-Hochberg technique, controlling for SES. Then, we evaluated statistical differences between envirotypes at the level of brain systems accounting for the spatial embedding and contiguity of the brain and using a ‘spin’ test. Briefly, this involved computing the mean difference in FC between every pair of brain systems and comparing this value against a null distribution generated using a space-preserving null model. FC differences were statistically significant if the observed difference exceeded that of the null distribution (false discovery rate fixed at *q* = 0.05; adjusted critical value of *p_crit_* = 0.02).

Then, we assessed the extent to which the pattern of brain network connectivity associated with better mental health is distinguishable for different dynamic social envirotypes. Specifically, we ran linear regressions predicting mental health outcomes (from Year 2) from dynamic social envirotype (informed by all 4 years) and a given edge weight (recorded at baseline) controlling for SES. This procedure involved one regression per functional connectivity edge weight, so we corrected for multiple comparisons. For these regressions, we were interested in whether envirotype interacted significantly with functional connectivity. After obtaining the *t*-statistics for the interaction terms, we accounted for the spatial embedding and contiguity of the brain by using the ‘spin’ test procedure described above. This process yielded both *t*-statistic matrices for each envirotype at the level of edge weights indicating how connectivity is associated with mental health outcomes for each envirotype and *Z*-statistic matrices for each envirotype pair at the level of edge weights.

## Supplementary Materials

### MRI Acquisition and Pre-processing

After completing motion compliance training in a simulated scanning environment, subjects first underwent a structural T1-weighted scan. Then, subjects completed two five-minute resting-state scans, during which they were instructed to lay with their eyes open while a crosshair was on the screen. After the two resting-state scans, subjects completed two other structural scans as part of the larger ABCD protocol, followed by one or two more resting-state scans, depending on the protocol at the specific study site. All scans were collected on one of three 3T scanner platforms with an adult-size head coil.

The functional MRI data we used were minimally pre-processed according to the HCP minimal pre-processing pipeline described in Glasser et al. (2013). Briefly, this pipeline includes distortion correction and alignment, denoising with Advanced Normalization Tools (ANTS89), FreeSurfer90 segmentation, surface registration, and volume registration using FSL FLIRT rigid-body transformation. Additional processing was done according to the DCAN BOLD Processing (DBP; Feczko et al. (2021)) pipeline which included the following steps: (1) DBP standard pre-processing, (2) removing respiratory signal from motion realignment data by filtering out frequencies between 18.582 - s25.726 breaths per minute, (3) applying DPB motion censoring (frames exceeding an FD threshold of 0.2mm or failing to pass outlier detection at *±*0.3 standard deviations were discarded), and (4) DBP generation of parcellated time series into the Gordon 333 cortical and 19 subcortical atlas Gordon et al. (2017).

### Modularity Maximization Algorithm

In general, the modularity, *Q*, of a partition can be expressed as the sum of contributions made by each community, *c ∈* 1*, …, K*, such that:

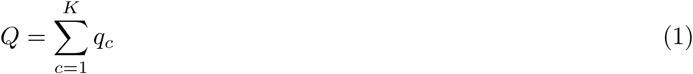

where

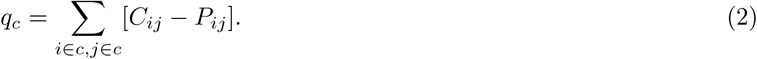

In this expression, *i* and *j* correspond to distinct elements in the matrix (i.e., distinct participants). The values of *C_ij_* and *P_ij_* correspond to the observed and expected correlation between those pairs of subjects, respectively. For a given correlation matrix, we uniformly set the expected weight of connections equal to the mean correlation value (i.e., we use a uniform null model). That is *P_ij_* = *C̅_ij_* for all *i, j* pairs, where 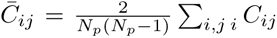 and *N_p_* is the total number of participants.

### Validating Clusters

We validated the cluster composition using two different techniques. First, we ensured the observed clusters were a good fit to the data by running the clustering algorithm (which is stochastic) 1000 times and computing the co-assignment matrix (i.e., the fraction of runs for which two subjects were assigned to the same cluster). We then identified consensus clusters with this co-assignment matrix, which are the clusters we report in the main text. To assess the extent to which clusters were meaningfully different from one another, we used a shuffling procedure 1000 times. We shuffled the rows and columns of the matrix defined by each subject’s score on each of the social environment measures, thereby breaking any associations across measures or people. For each shuffled matrix, we computed the pairwise correlation between all people across all measures, performed community detection, and calculated the modularity *Q* of the optimal partition. *Q* is larger when people in the same community are more similar than people in different communities. We compared the *Q*s resulting from the shuffled matrices to the value of *Q* from the empirical data.

---

**Table S1:** Demographic make-up of analyzed sample versus full ABCD and excluded sample. Percentages do not include participants who have missing data for these measures.

| Sample | Sex | Race | Ethnicity | Parent Education | Household Income |
| --- | --- | --- | --- | --- | --- |
| Full | 47.8% F | 4.4% Mixed, 79% White,<br>8.1% African American,<br>0.8% Native American,<br>0.1% Pacific Islander<br>2.8% Asian, 4.6% Other | 16.9% Hispanic or<br>Latino, 83.1% Not<br>Hispanic/Latino | 4.4% Some high school,<br>1.6% High school diploma<br>25.9% Some college,<br>5.7% College degree,<br>60.3% Advanced degree | 13.7% \$0 - \$24,999,<br>13.4% \$25,000 - \$49,999,<br>12.6% \$50,000 - \$74,999,<br>13.2% \$75,000 - \$99,999,<br>27.9% \$100,000 - \$199,999,<br>10.5% \$200,000 and up |
| Included | 47% F | 4.9% Mixed, 79.2% White,<br>7.9% African American,<br>0.8% Native American,<br>0.1% Pacific Islander,<br>2.9% Asian, 4% Other | 15.2% Hispanic or<br>Latino, 84.2% Not<br>Hispanic/Latino | 3.3% Some high school,<br>1.4% High school diploma,<br>0.02% Some college,<br>7.3% College degree,<br>65% Advanced degree | 10.7% \$0 - \$24,999,<br>4.9% \$25,000 - \$34,999,<br>7.8% \$35,000 - \$49,999,<br>13.9% \$50,000 - \$74,999,<br>15% \$75,000 - \$99,999,<br>34% \$100,000 - \$199,999,<br>13.4% \$200,000 and up |
| Excluded | 48.1% F | 2.8% Mixed, 78.7% White,<br>8.9% African American,<br>0.5% Native American,<br>0.2% Pacific Islander,<br>2.6% Asian, 6.2% Other | 17.8% Hispanic or<br>Latino, 82.2% Not<br>Hispanic/Latino | 5.1% Some high school,<br>1.8% High school diploma,<br>0.01% Some college,<br>8% College degree,<br>57.3% Advanced degree | 15.2% \$0 - \$24,999,<br>6.5% \$25,000 - \$34,999,<br>8.2% \$35,000 - \$49,999,<br>12.5% \$50,000 - \$74,999,<br>13.4% \$75,000 - \$99,999,<br>31.1% \$100,000 - \$199,999,<br>12.8% \$200,000 and up |

**Figure S1:**
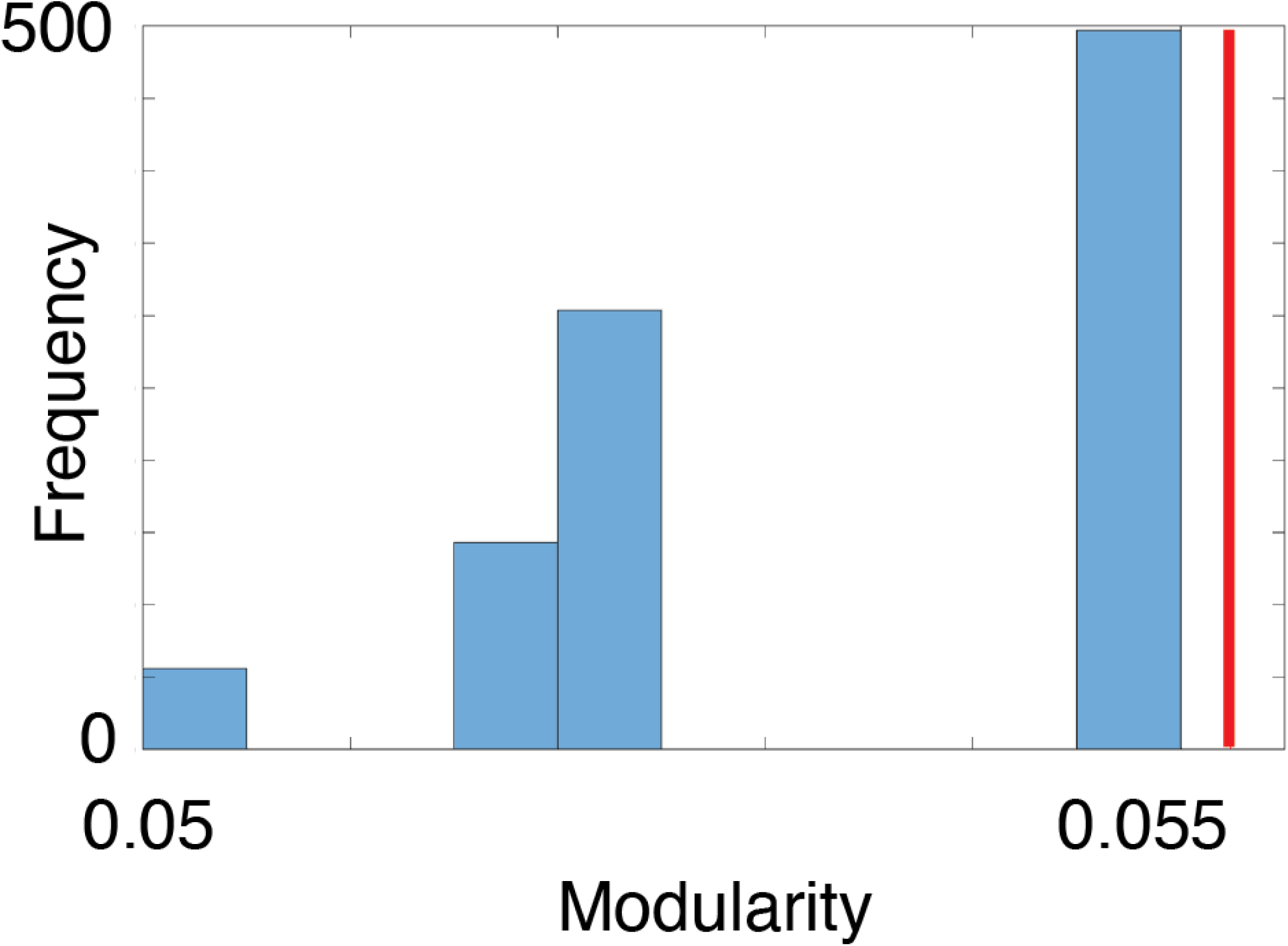
Validation of clusters using modularity. We compared the modularity of the observed clusters (red dashed line) with the modularity of 250 shuffled clusters (histogram). The higher modularity of the observed clusters than all other shuffled clusters indicates they are more internally similar and externally dissimilar than would be expected by chance. That is, these particular dynamic social envirotypes are sufficiently representative of the dominant modes of social environment change in the data.

**Figure S2:**
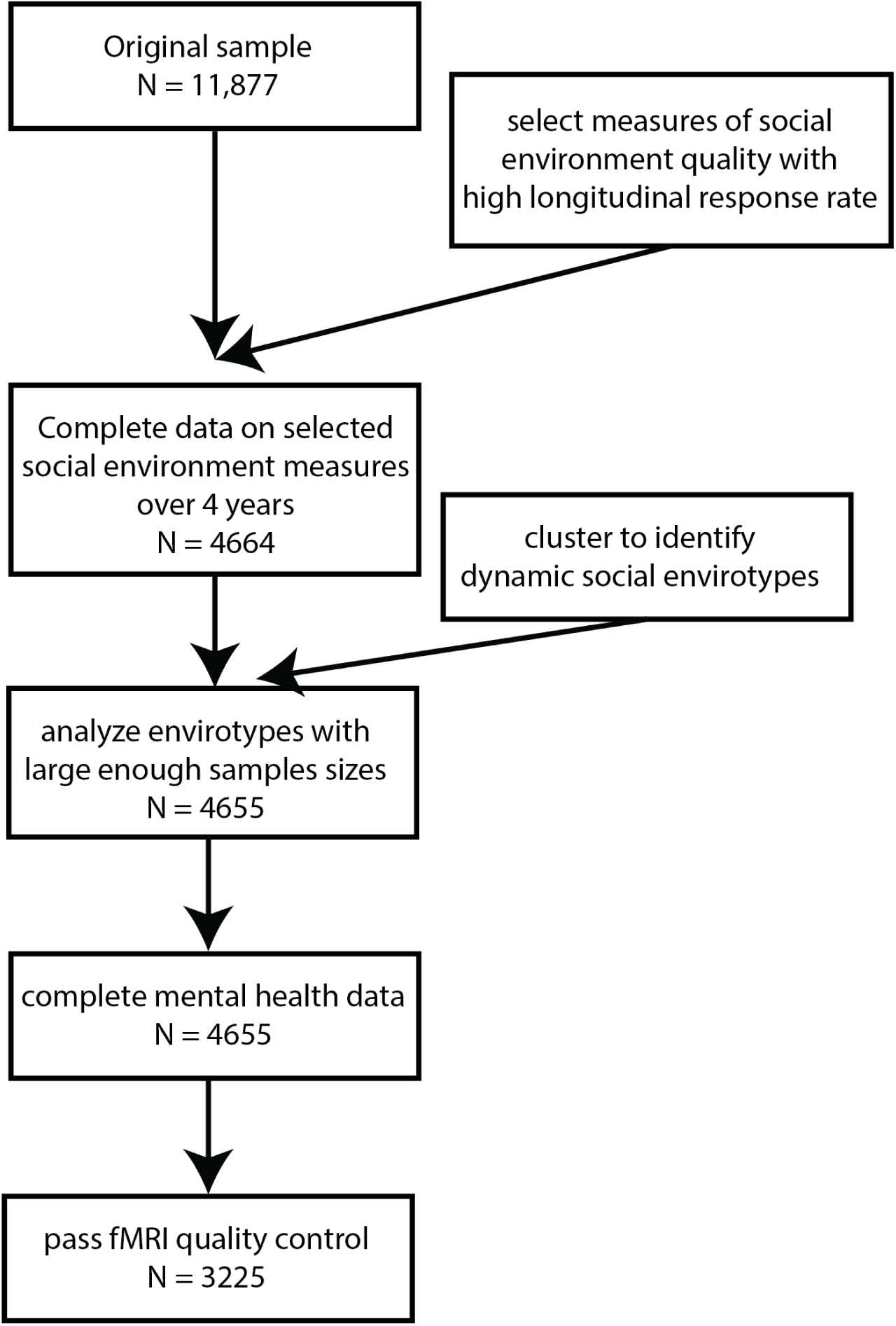
Inclusion tree. We show the sample size at each inclusion step.

**Table S2:**
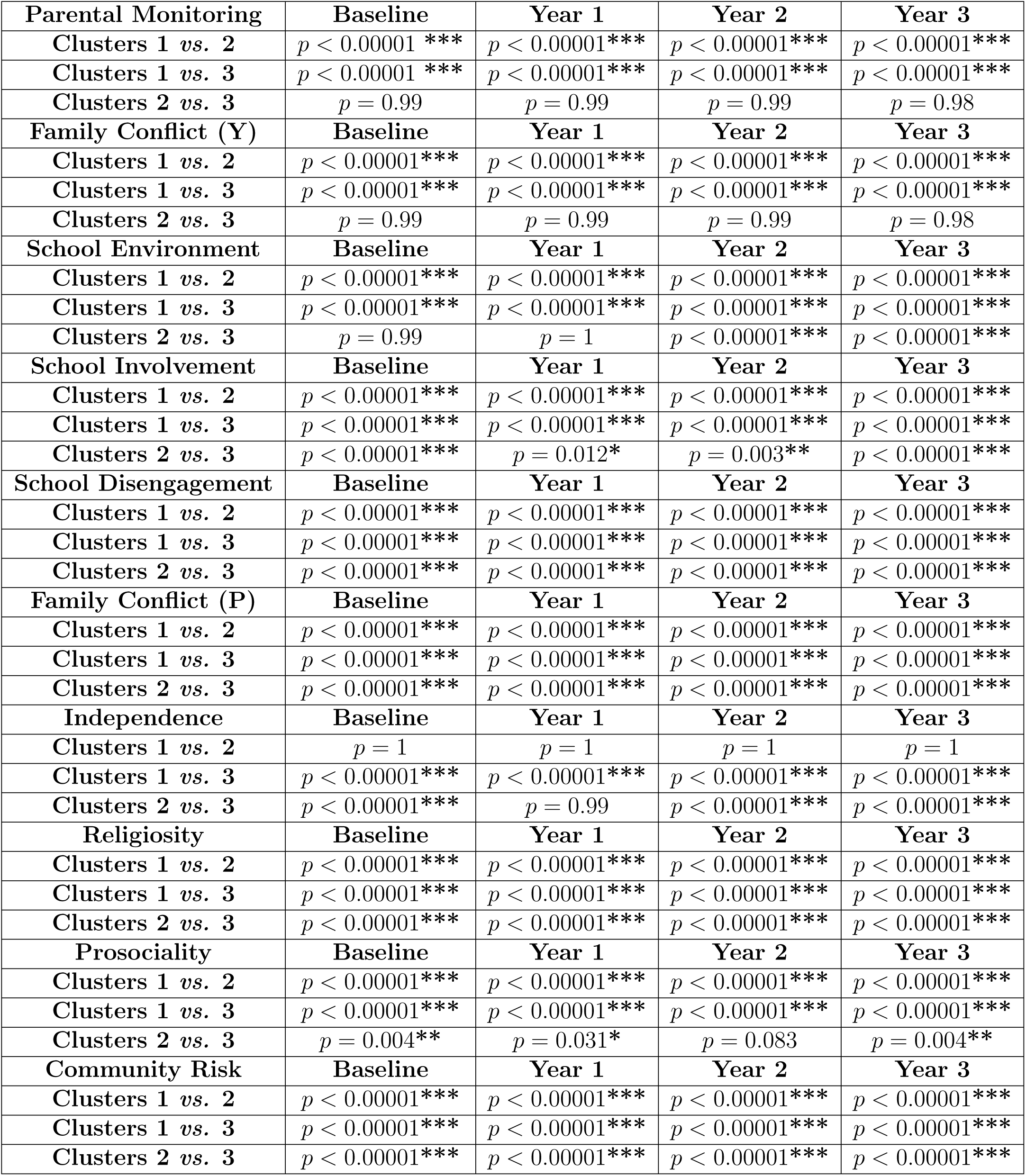
Adjusted *p*-values of cluster comparisons. *:*p <* 0.05, **:*p <* 0.01, ***:*p <* 0.001.

**Figure S3:**
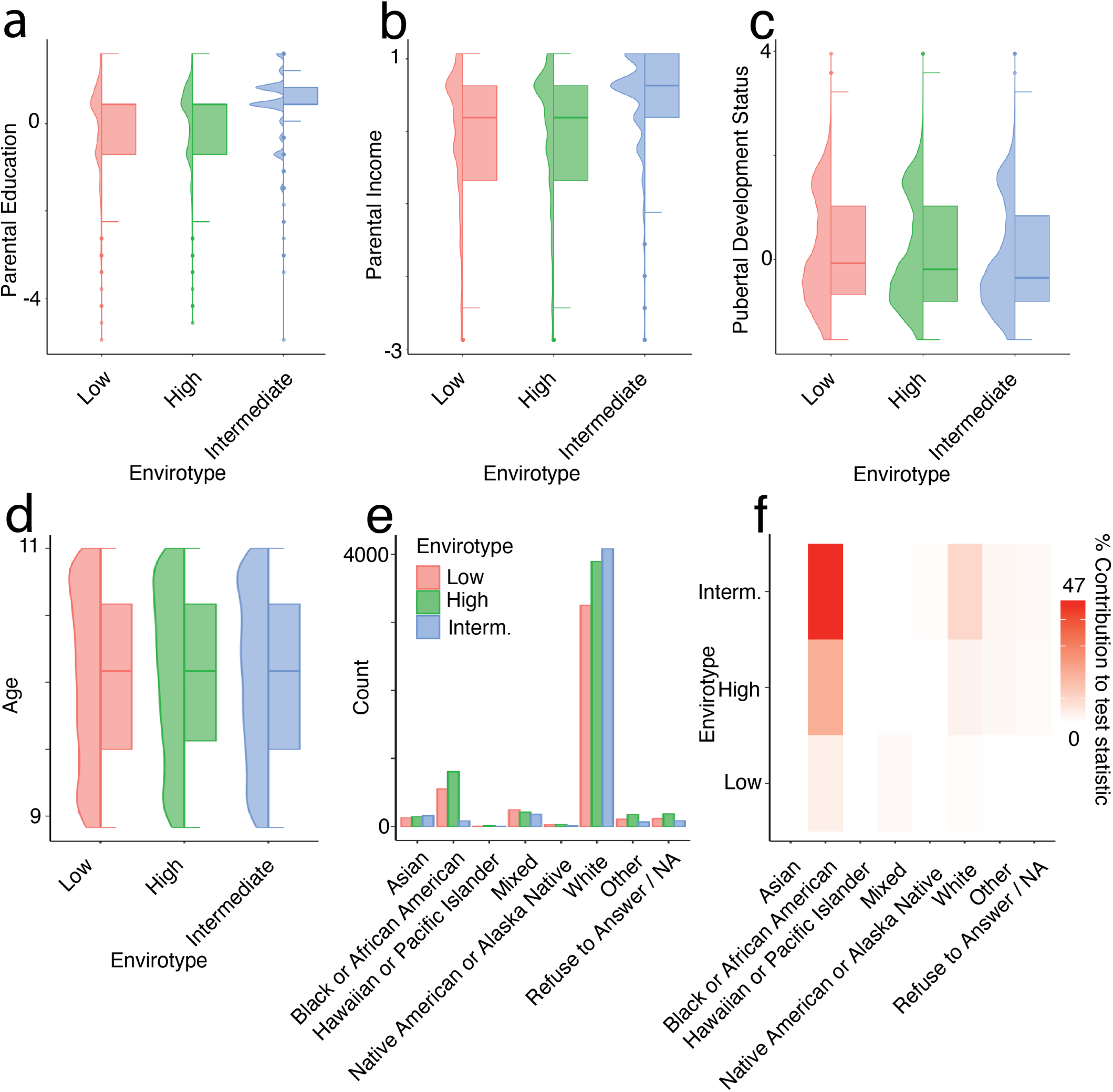
Cluster demographics. We assessed the composition of the clusters based on parental education, parental income, pubertal development status, and race/ethnicity.

**Figure S4:**
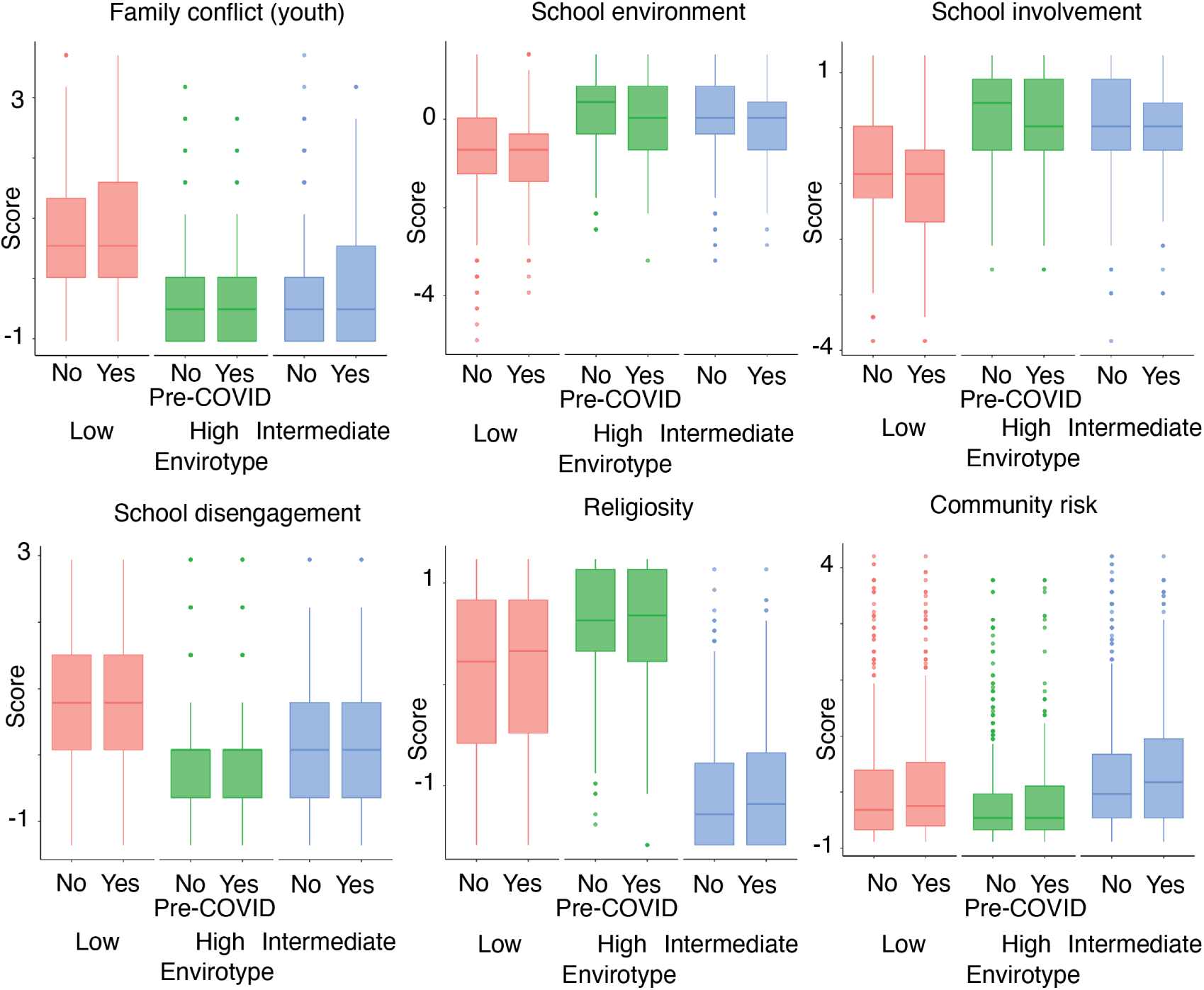
COVID timing across clusters. We examined how the COVID-19 lockdowns may have affected scores on the social environment measures. While there were significant main effects of COVID timing on six out of ten social environment measures (shown here), there were no inter-actions between COVID timing and cluster affiliation, meaning all clusters were similarly impacted by COVID.

**Figure S5:**
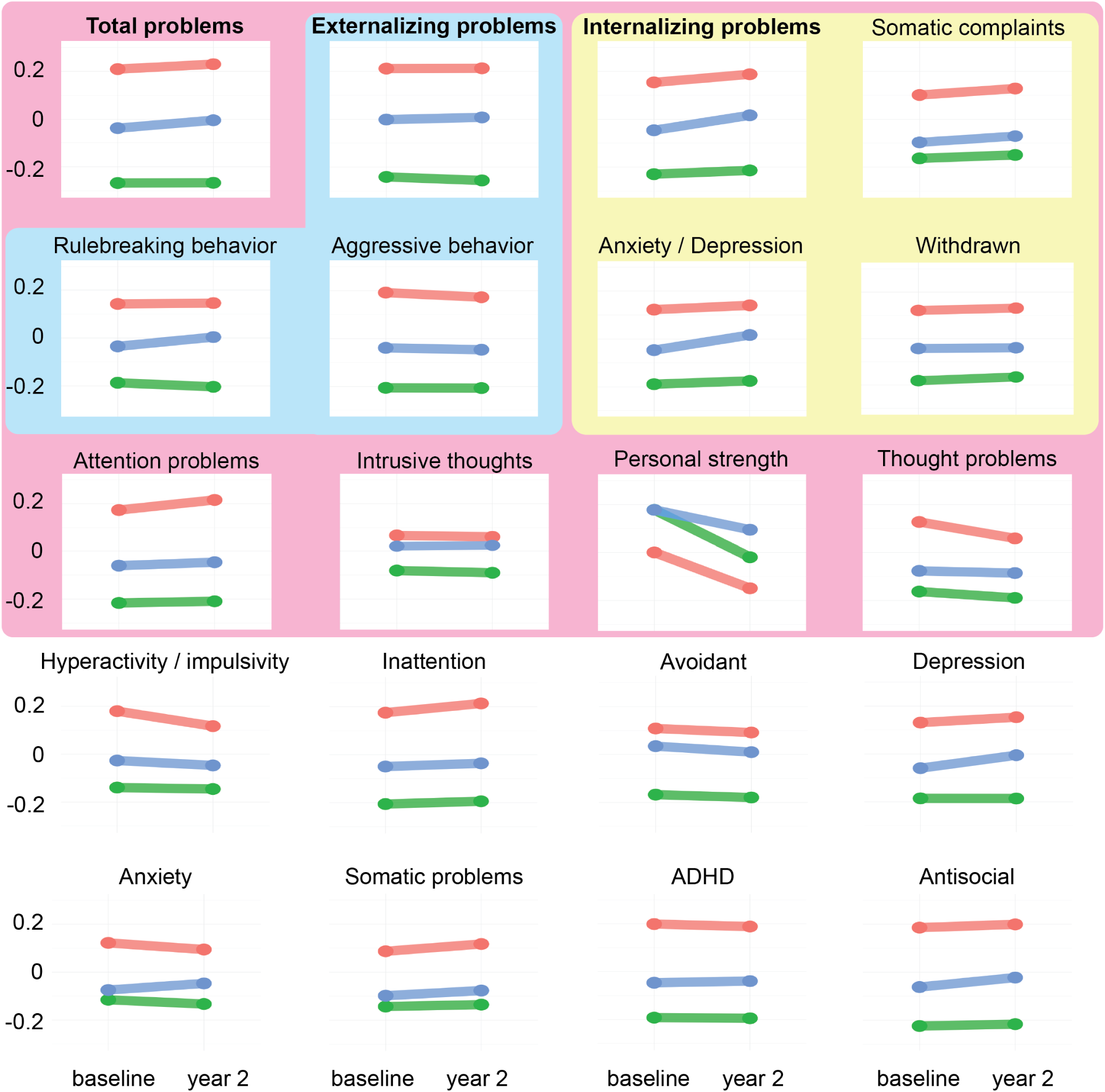
Cluster mental health. We looked at the mental health of the clusters at two time points, with the y-axis of all plots representing Z-scored prevalence of symptoms, so a higher score indicates worse mental health for all measures except Personal Strength. Similarly, a negative (positive) slope indicates improvement (worsening) in mental health. Across the three clusters, mental health starts and ends differently. Composite measures are indicated by bold text, and the component measures used to compute them are indicated by the boxes.

**Figure S6:**
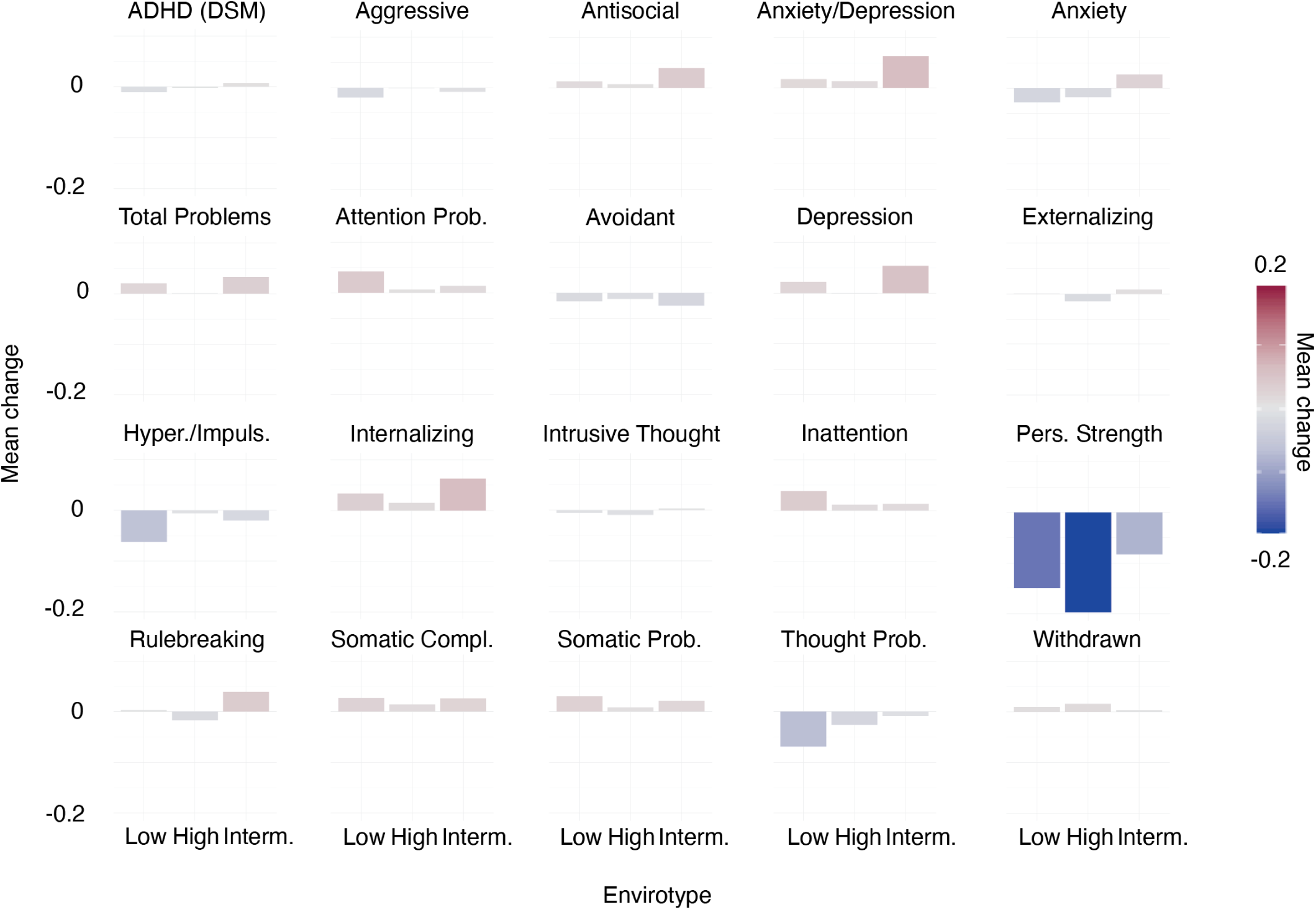
Changes in cluster mental health.

**Figure S7:**
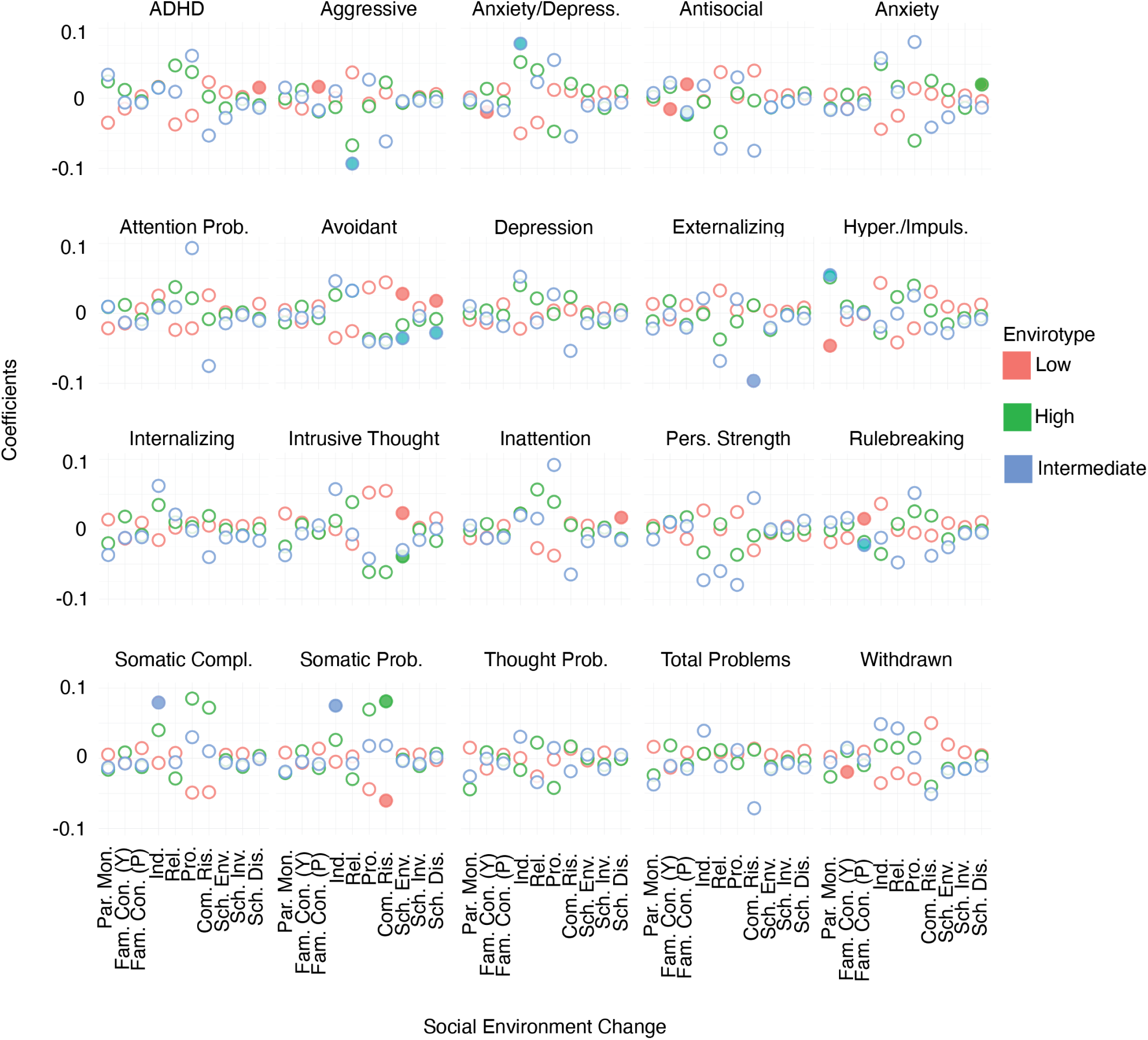
Linking change in the social environment to change in mental health. We assessed whether changes in the social environment measures were associated with outcomes in mental health measures recorded in year 2. No comparisons survived multiple comparisons corrections, but we show here uncorrected test statistics. Significant comparisons before correction are indicated by filled circles.

**Figure S8:**
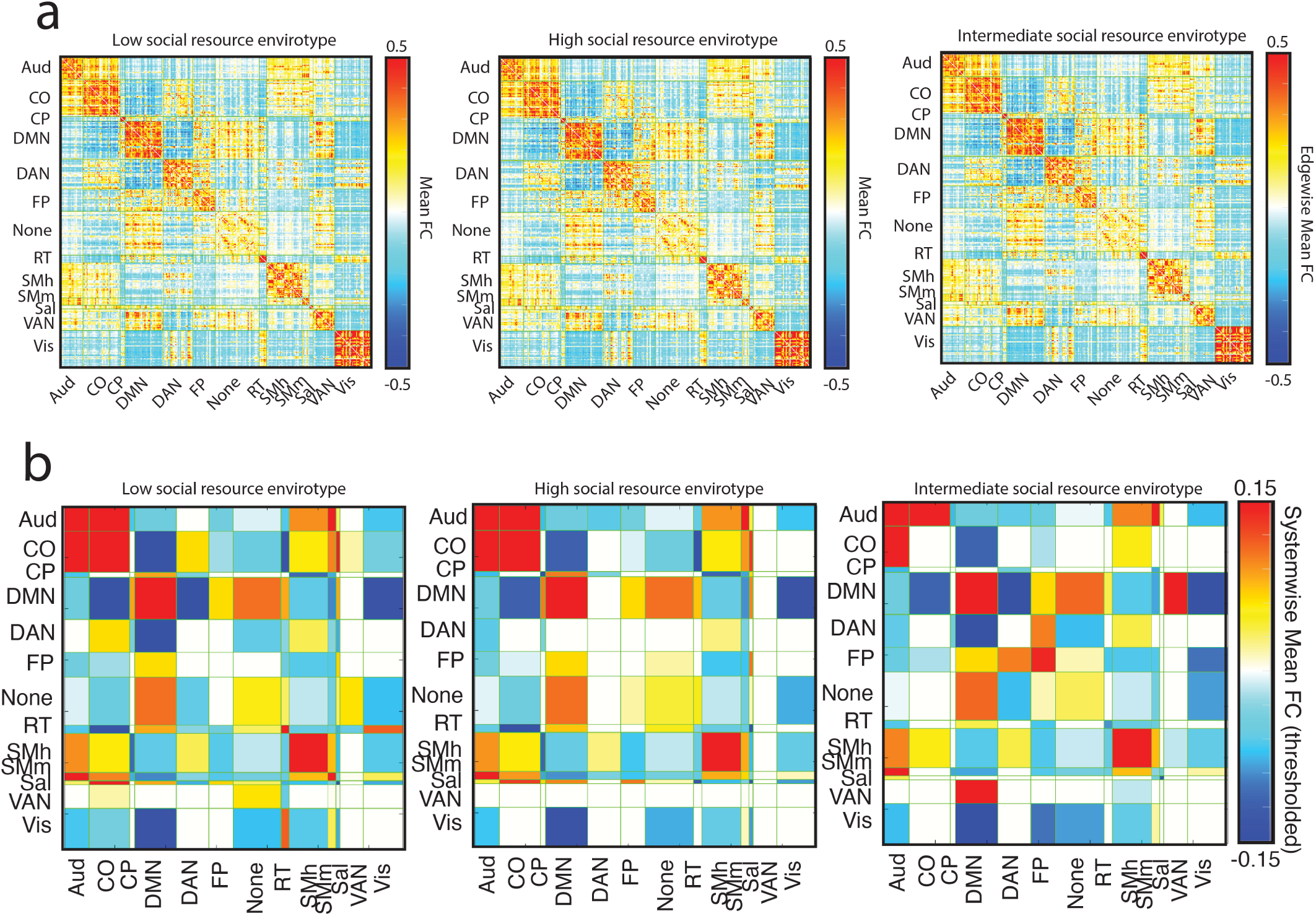
Characteristic brain network organization for each dynamic social envirotype. Row (a) displays the mean functional connectivity between systems at the node level for each cluster. Brighter red (or darker blue) indicates that a cluster tends to have higher (or lower) connectivity between that pair of systems. Row (b) summarizes the characteristic at the system level, thresholded to display differences that are statistically significant.

**Figure S9:**
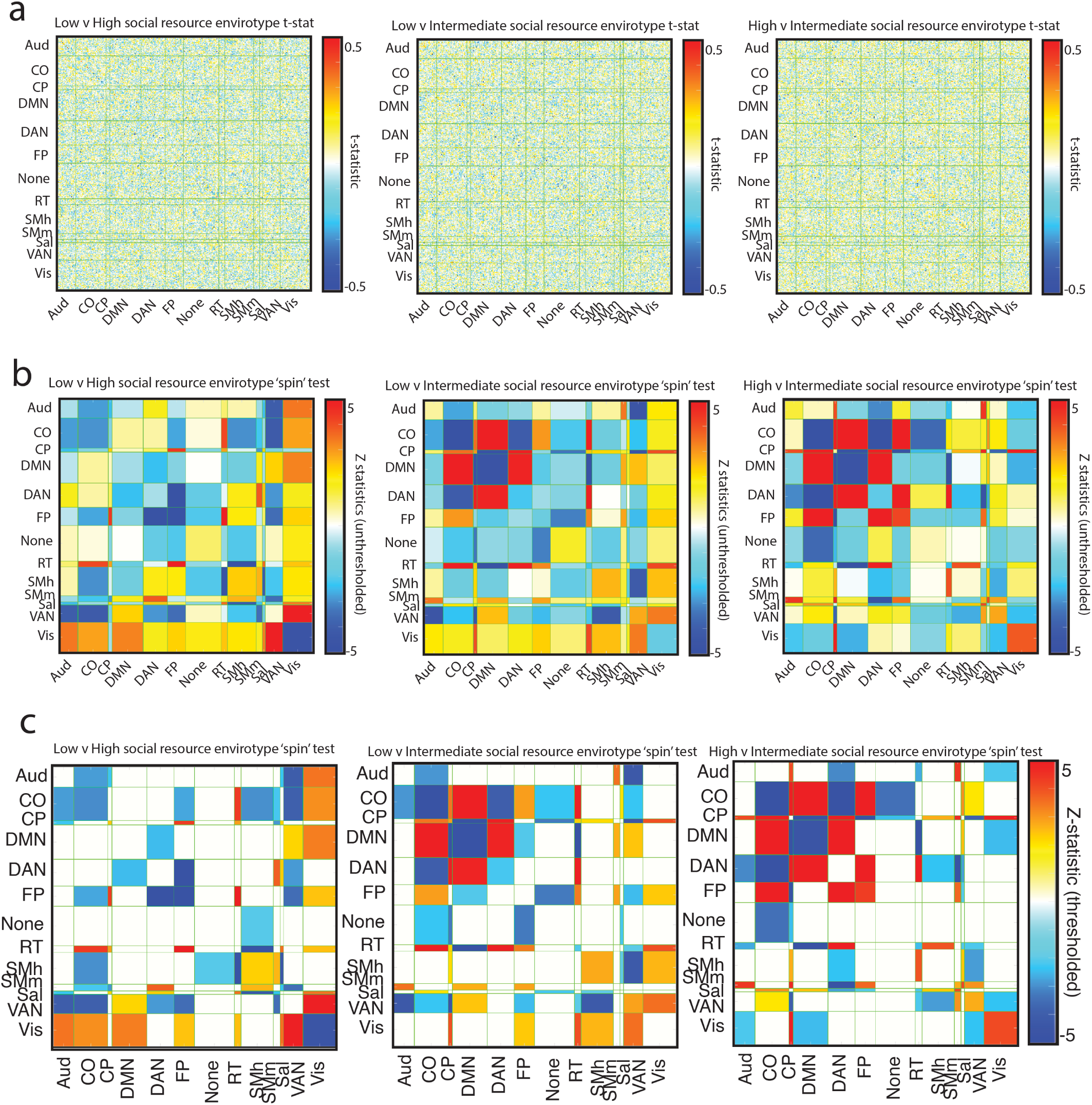
Brain networks are organized differently across different dynamic social envirotypes. Row (a) shows the unthresholded *t*-statistic matrix for clusterwise comparisons. Row(b) reflects the unthresholded *Z*-statistics from the ‘spin’ tests, the thresholded (based on statistical significance) versions of which are shown in row (c). Brighter red (or darker blue) means that cluster *i* has significantly higher (lower) connectivity than cluster *j* between that pair of systems.

**Figure S10:**
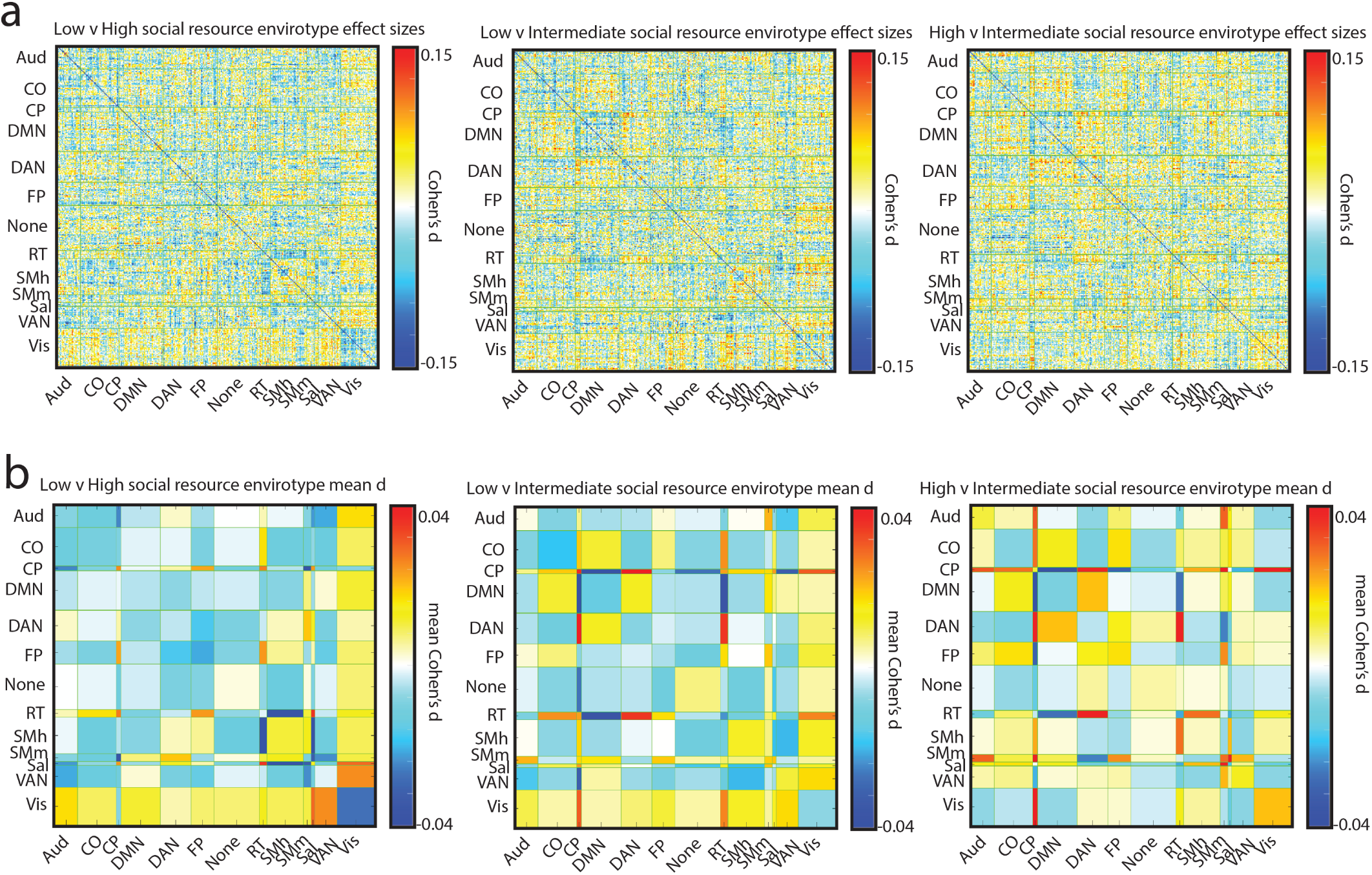
Effect sizes for differences in brain network connectivity across clusters. Row (a) shows Cohen’s *d* for all pairwise comparisons at the edge level. Row (b) depicts the mean Cohen’s *d* at the system level.

**Figure S11:**
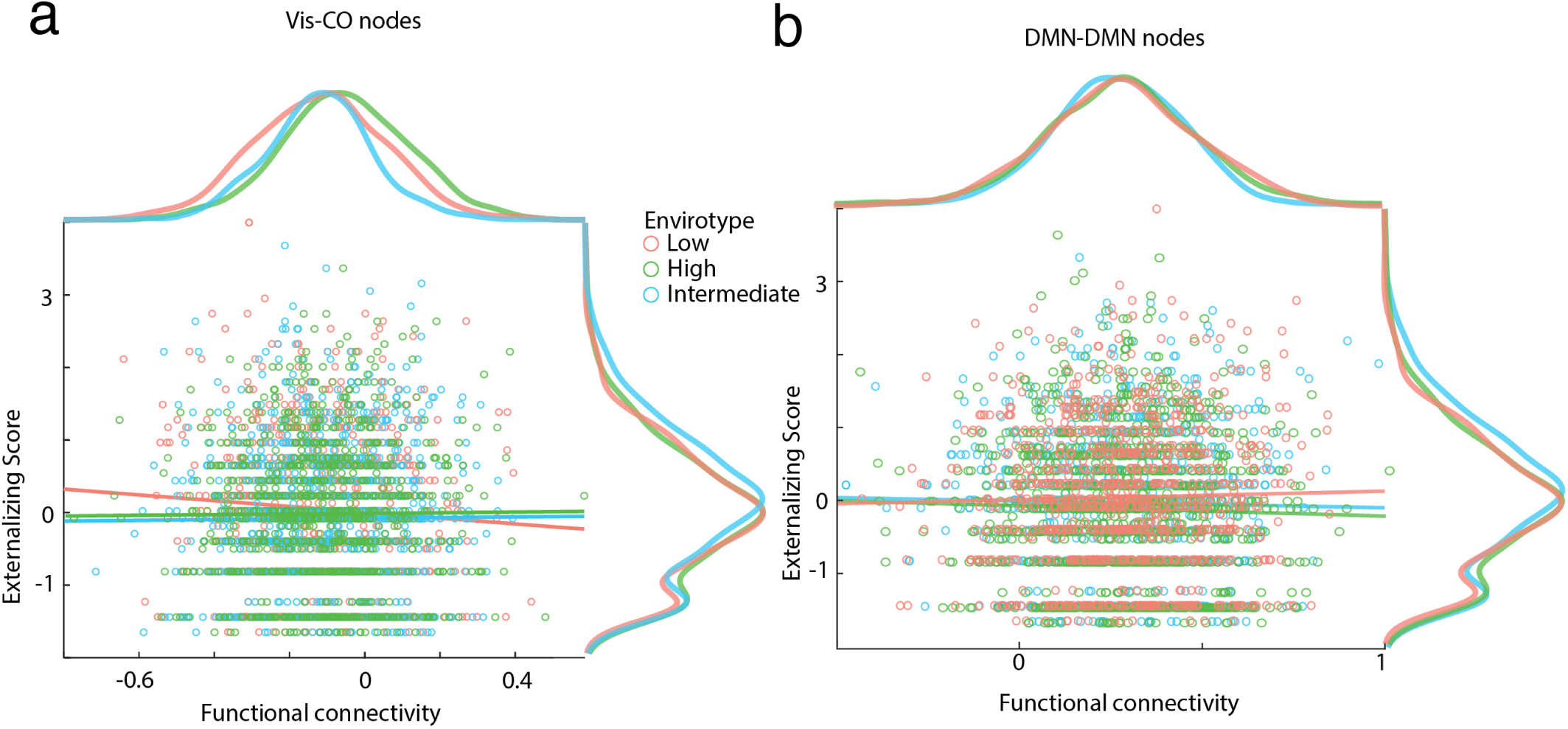
Scatterplots of functional connectivity and externalizing problems by cluster. Panels (a) and (b) show the relationship between Vis-CO and intra-DMN, respectively, functional connectivity and externalizing problems by cluster. The *t*-statistics for these associations across the whole brain are shown in Supplementary Figure S12, and the thresholded, *Z*-statistics that summarise results at the system level are in Figure S13.

**Figure S12:**
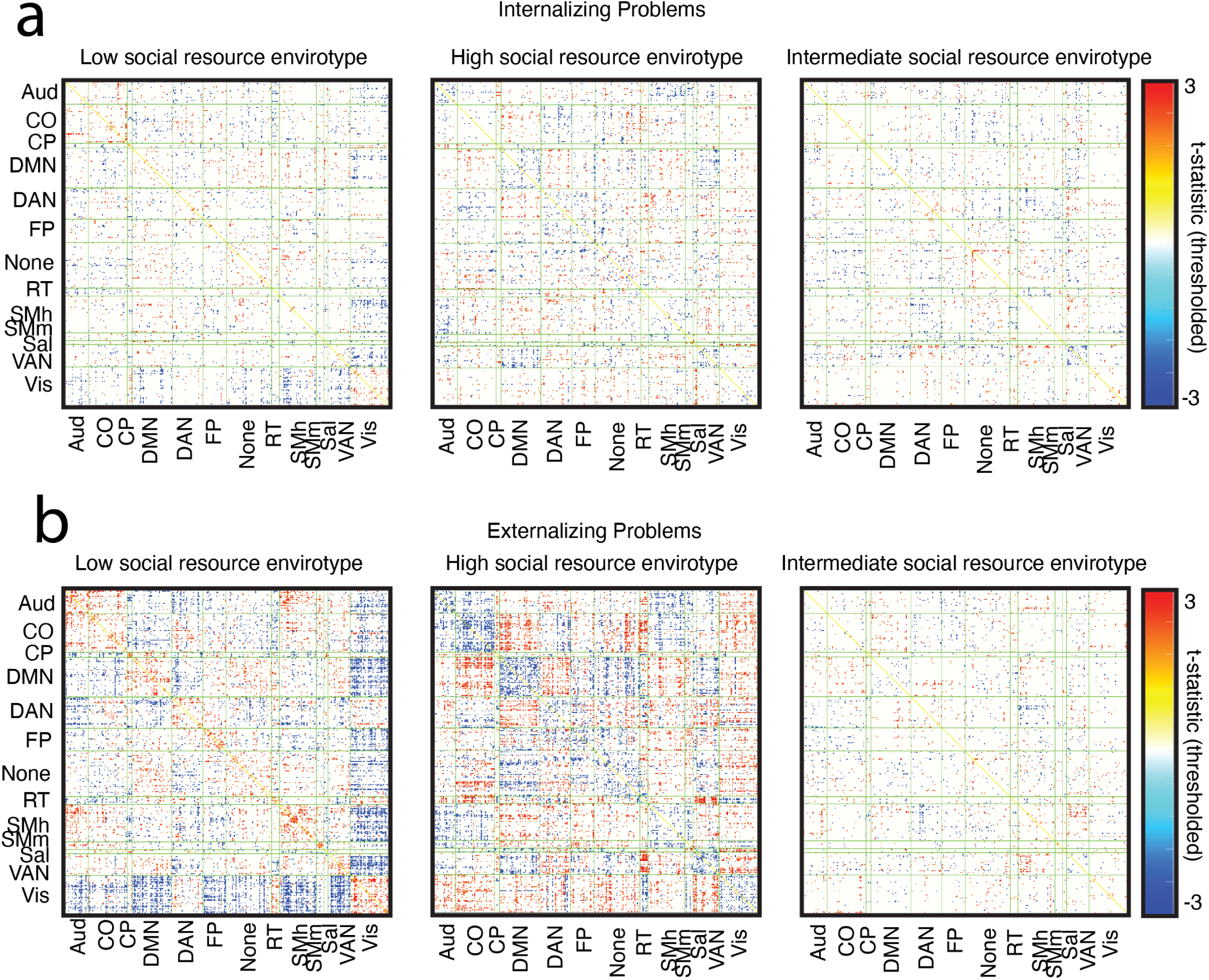
Thresholded t-statistic matrices for moderation analysis of FC differences across clusters. Row (a) shows thresholded *t*-statistic matrices for each cluster’s association between brain network functional connectivity and internalizing problems. Row (b) shows the same but for externalizing problems. All tests are corrected for multiple comparisons and thresholded according to the adjusted *p_crit_*= 0.0415.

**Figure S13:**
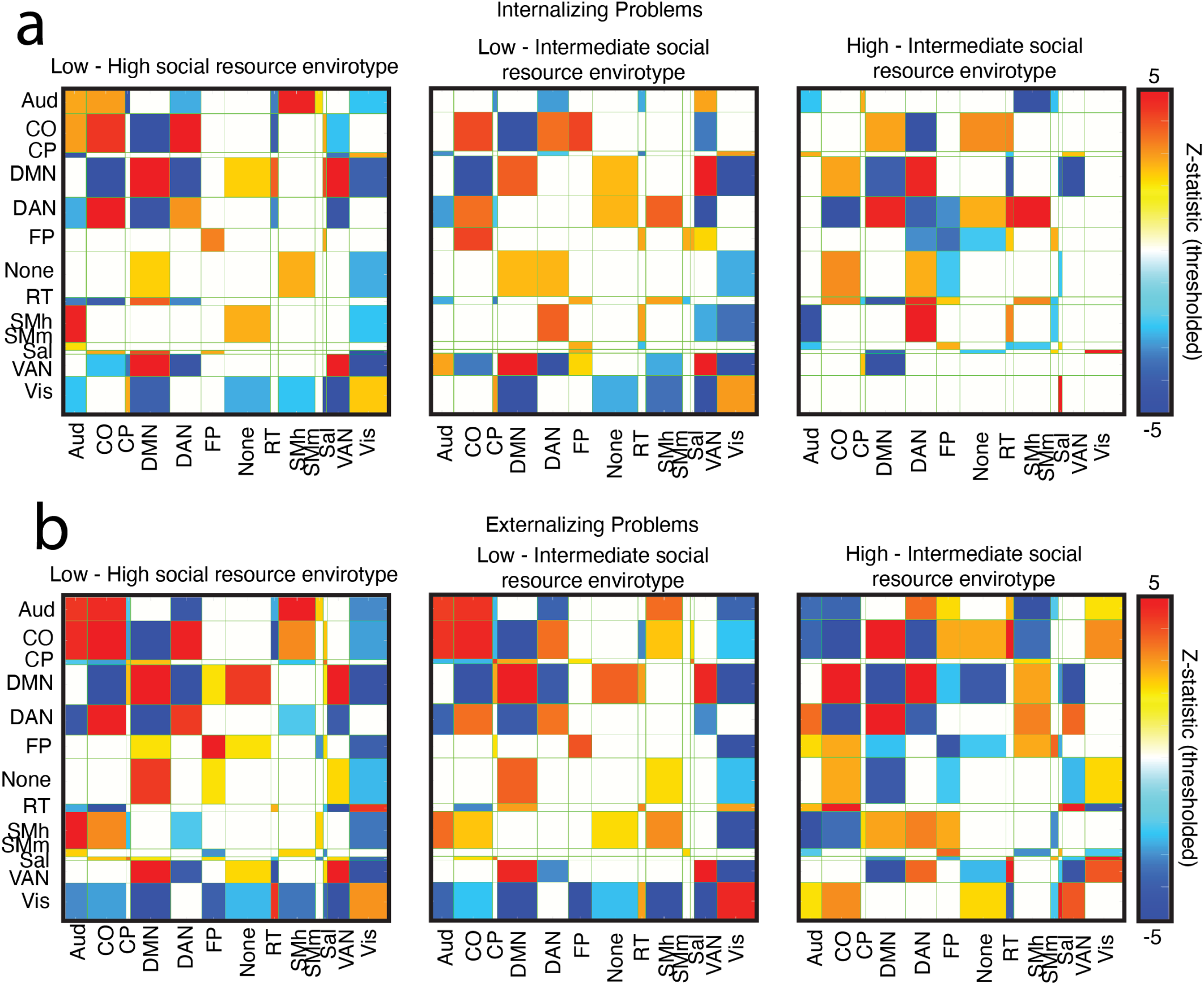
Differences in the network neural underpinnings of mental health. We assess the extent to which internalizing problems (row a) and externalizing problems (row b) are associated with distinguishable patterns of connectivity for the different clusters, aggregating at the system-level using space-preserving ‘spin’ tests. Both rows show thresholded *Z*-statistics of the association between mental health and system-wise connectivity for each cluster comparison.

**Figure S14:**
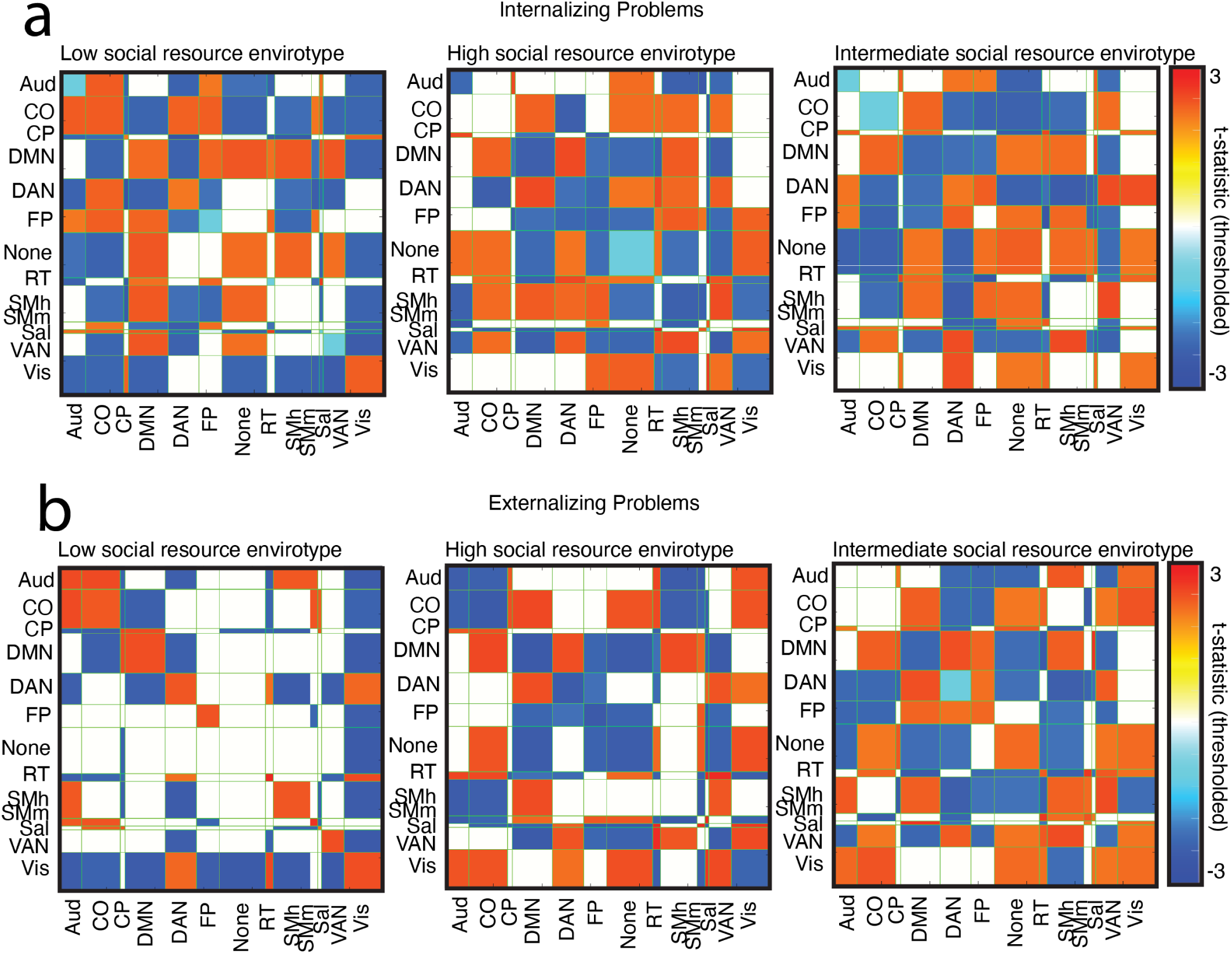
Brain networks associated with better mental health are organized differently across different dynamic social envirotypes. We assess the extent to which internalizing problems (row a) and externalizing problems (row b) are associated with distinguishable patterns of connectivity for the different clusters. In both cases, brighter red indicates a positive association (higher connectivity is associated with more symptoms), while darker blue indicates a negative association (higher connectivity is associated with fewer symptoms).

**Table S3:** Adjusted *p*-values of within-cluster temporal comparisons. *:*p <* 0.05, **:*p <* 0.01, ***:p < 0.001.

|  |  |  |  |
| --- | --- | --- | --- |
| **Parental Monitoring** | Baseline vs. Year 1 | Year 1 vs. Year 2 | Year 2 vs. Year 3 |
| Cluster 1 | $p < 0.00001^{***}$ | $p = 0.99$ | $p < 0.00001^{***}$ |
| Cluster 2 | $p < 0.00001^{***}$ | $p = 1$ | $p < 0.00001^{***}$ |
| Cluster 3 | $p < 0.00001^{***}$ | $p = 1$ | $p < 0.00001^{***}$ |
| **Family Conflict (Youth)** | Baseline vs. Year 1 | Year 1 vs. Year 2 | Year 2 vs. Year 3 |
| Cluster 1 | $p = 0.99$ | $p = 1$ | $p = 0.46$ |
| Cluster 2 | $p = 0.1$ | $p = 0.99$ | $p = 0.95$ |
| Cluster 3 | $p = 0.00001^{***}$ | $p = 1$ | $p < 0.00001^{***}$ |
| **School Environment** | Baseline vs. Year 1 | Year 1 vs. Year 2 | Year 2 vs. Year 3 |
| Cluster 1 | $p = 1$ | $p = 0.99$ | $p < 0.00001^{***}$ |
| Cluster 2 | $p < 0.00001^{***}$ | $p = 1$ | $p = 0.99$ |
| Cluster 3 | $p < 0.00001^{***}$ | $p = 1$ | $p < 0.00001^{***}$ |
| **School Involvement** | Baseline vs. Year 1 | Year 1 vs. Year 2 | Year 2 vs. Year 3 |
| Cluster 1 | $p = 1$ | $p < 0.00001^{***}$ | $p = 1$ |
| Cluster 2 | $p = 0.003^{**}$ | $p < 0.00001^{***}$ | $p = 0.56$ |
| Cluster 3 | $p < 0.00001^{***}$ | $p < 0.00001^{***}$ | $p = 0.018^*$ |
| **School Disengagement** | Baseline vs. Year 1 | Year 1 vs. Year 2 | Year 2 vs. Year 3 |
| Cluster 1 | $p = 1$ | $p = 0.0006^{***}$ | $p = 0.604$ |
| Cluster 2 | $p = 0.5$ | $p < 0.00001^{***}$ | $p < 0.00001^{***}$ |
| Cluster 3 | $p = 1$ | $p = 0.001^{**}$ | $p < 0.00001^{***}$ |
| **Family Conflict (Parent)** | Baseline vs. Year 1 | Year 1 vs. Year 2 | Year 2 vs. Year 3 |
| Cluster 1 | $p = 1$ | $p = 1$ | $p = 1$ |
| Cluster 2 | $p = 0.84$ | $p = 1$ | $p = 0.99$ |
| Cluster 3 | $p = 1$ | $p = 1$ | $p = 1$ |
| **Independence** | Baseline vs. Year 1 | Year 1 vs. Year 2 | Year 2 vs. Year 3 |
| Cluster 1 | $p = 0.88$ | $p = 1$ | $p = 0.0002^{***}$ |
| Cluster 2 | $p = 0.82$ | $p = 0.72$ | $p = 0.001^{**}$ |
| Cluster 3 | $p < 0.00001^{***}$ | $p = 0.78$ | $p = 0.41$ |
| **Religiosity** | Baseline vs. Year 1 | Year 1 vs. Year 2 | Year 2 vs. Year 3 |
| Cluster 1 | $p = 0.42$ | $p = 1$ | $p = 0.016^*$ |
| Cluster 2 | $p = 1$ | $p = 1$ | $p = 0.0009^{***}$ |
| Cluster 3 | $p = 0.5$ | $p = 1$ | $p = 1$ |
| **Prosociality** | Baseline vs. Year 1 | Year 1 vs. Year 2 | Year 2 vs. Year 3 |
| Cluster 1 | $p = 0.036^*$ | $p = 1$ | $p = 0.003^{**}$ |
| Cluster 2 | $p = 1$ | $p = 1$ | $p = 0.48$ |
| Cluster 3 | $p = 1$ | $p = 1$ | $p = 0.216$ |
| **Community Risk** | Baseline vs. Year 1 | Year 1 vs. Year 2 | Year 2 vs. Year 3 |
| Cluster 1 | $p = 1$ | $p = 0.091$ | $p = 1$ |
| Cluster 2 | $p = 1$ | $p = 0.073$ | $p = 0.99$ |
| Cluster 3 | $p < 0.00001^{***}$ | $p = 1$ | $p < 0.00001^{***}$ |

**Table S4:** Effect sizes for cluster-wise comparisons of scores on social measures.

| Measure | Year | Cluster | Year:Cluster |
| --- | --- | --- | --- |
| Parental Monitoring | $\eta^2 = 0.01$ | $\eta^2 = 0.15$ | $\eta^2 = 1.09 \times 10^{-4}$ |
| Family Conflict (Youth) | $\eta^2 = 1.88 \times 10^{-3}$ | $\eta^2 = 0.17$ | $\eta^2 = 1.01 \times 10^{-3}$ |
| Family Conflict (Parent) | $\eta^2 = 4.42 \times 10^{-4}$ | $\eta^2 = 0.08$ | $\eta^2 = 4.69 \times 10^{-4}$ |
| School Environment | $\eta^2 = 0.02$ | $\eta^2 = 0.16$ | $\eta^2 = 1.22 \times 10^{-3}$ |
| School Involvement | $\eta^2 = 0.02$ | $\eta^2 = 0.23$ | $\eta^2 = 7.34 \times 10^{-4}$ |
| School Disengagement | $\eta^2 = 0.02$ | $\eta^2 = 0.16$ | $\eta^2 = 6.06 \times 10^{-4}$ |
| Independence | $\eta^2 = 4.71 \times 10^{-3}$ | $\eta^2 = 0.01$ | $\eta^2 = 6.17 \times 10^{-4}$ |
| Religiosity | $\eta^2 = 1.85 \times 10^{-3}$ | $\eta^2 = 0.56$ | $\eta^2 = 4.91 \times 10^{-4}$ |
| Prosociality | $\eta^2 = 3.11 \times 10^{-3}$ | $\eta^2 = 0.07$ | $\eta^2 = 7.33 \times 10^{-4}$ |
| Community Risk | $\eta^2 = 2.12 \times 10^{-3}$ | $\eta^2 = 0.05$ | $\eta^2 = 3.86 \times 10^{-4}$ |

**Table S5:**
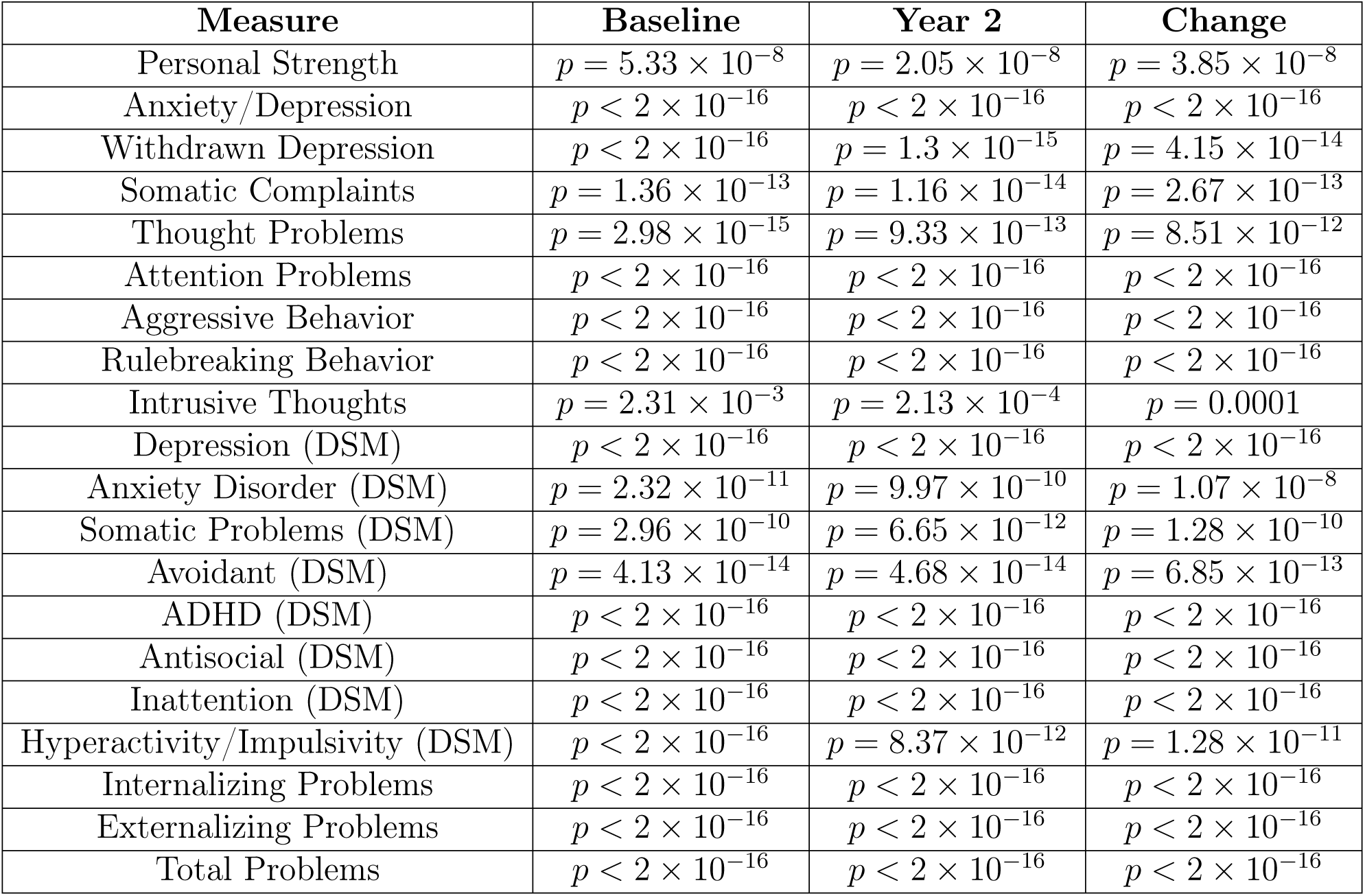
Adjusted *p*-values for omnibus cluster-wise comparisons of mental health scores.

| Measure | Baseline | Year 2 | Change |
| --- | --- | --- | --- |
| Personal Strength | $p = 5.33 \times 10^{-8}$ | $p = 2.05 \times 10^{-8}$ | $p = 3.85 \times 10^{-8}$ |
| Anxiety/Depression | $p < 2 \times 10^{-16}$ | $p < 2 \times 10^{-16}$ | $p < 2 \times 10^{-16}$ |
| Withdrawn Depression | $p < 2 \times 10^{-16}$ | $p = 1.3 \times 10^{-15}$ | $p = 4.15 \times 10^{-14}$ |
| Somatic Complaints | $p = 1.36 \times 10^{-13}$ | $p = 1.16 \times 10^{-14}$ | $p = 2.67 \times 10^{-13}$ |
| Thought Problems | $p = 2.98 \times 10^{-15}$ | $p = 9.33 \times 10^{-13}$ | $p = 8.51 \times 10^{-12}$ |
| Attention Problems | $p < 2 \times 10^{-16}$ | $p < 2 \times 10^{-16}$ | $p < 2 \times 10^{-16}$ |
| Aggressive Behavior | $p < 2 \times 10^{-16}$ | $p < 2 \times 10^{-16}$ | $p < 2 \times 10^{-16}$ |
| Rulebreaking Behavior | $p < 2 \times 10^{-16}$ | $p < 2 \times 10^{-16}$ | $p < 2 \times 10^{-16}$ |
| Intrusive Thoughts | $p = 2.31 \times 10^{-3}$ | $p = 2.13 \times 10^{-4}$ | $p = 0.0001$ |
| Depression (DSM) | $p < 2 \times 10^{-16}$ | $p < 2 \times 10^{-16}$ | $p < 2 \times 10^{-16}$ |
| Anxiety Disorder (DSM) | $p = 2.32 \times 10^{-11}$ | $p = 9.97 \times 10^{-10}$ | $p = 1.07 \times 10^{-8}$ |
| Somatic Problems (DSM) | $p = 2.96 \times 10^{-10}$ | $p = 6.65 \times 10^{-12}$ | $p = 1.28 \times 10^{-10}$ |
| Avoidant (DSM) | $p = 4.13 \times 10^{-14}$ | $p = 4.68 \times 10^{-14}$ | $p = 6.85 \times 10^{-13}$ |
| ADHD (DSM) | $p < 2 \times 10^{-16}$ | $p < 2 \times 10^{-16}$ | $p < 2 \times 10^{-16}$ |
| Antisocial (DSM) | $p < 2 \times 10^{-16}$ | $p < 2 \times 10^{-16}$ | $p < 2 \times 10^{-16}$ |
| Inattention (DSM) | $p < 2 \times 10^{-16}$ | $p < 2 \times 10^{-16}$ | $p < 2 \times 10^{-16}$ |
| Hyperactivity/Impulsivity (DSM) | $p < 2 \times 10^{-16}$ | $p = 8.37 \times 10^{-12}$ | $p = 1.28 \times 10^{-11}$ |
| Internalizing Problems | $p < 2 \times 10^{-16}$ | $p < 2 \times 10^{-16}$ | $p < 2 \times 10^{-16}$ |
| Externalizing Problems | $p < 2 \times 10^{-16}$ | $p < 2 \times 10^{-16}$ | $p < 2 \times 10^{-16}$ |
| Total Problems | $p < 2 \times 10^{-16}$ | $p < 2 \times 10^{-16}$ | $p < 2 \times 10^{-16}$ |

**Table S6:** Effect sizes for cluster-wise comparisons of mental health scores.

| Measure | Baseline | Year 2 | Change |
| --- | --- | --- | --- |
| Personal Strength | $\eta^2 = 7.4 \times 10^{-3}$ | $\eta^2 = 7.72 \times 10^{-3}$ | $\eta^2 = 0.06$ |
| Anxiety/Depression | $\eta^2 = 0.02$ | $\eta^2 = 0.02$ | $\eta^2 = 0.02$ |
| Withdrawn Depression | $\eta^2 = 0.02$ | $\eta^2 = 0.02$ | $\eta^2 = 0.02$ |
| Somatic Complaints | $\eta^2 = 0.01$ | $\eta^2 = 7.72 \times 10^{-3}$ | $\eta^2 = 8.75 \times 10^{-3}$ |
| Thought Problems | $\eta^2 = 0.01$ | $\eta^2 = 0.01$ | $\eta^2 = 0.01$ |
| Attention Problems | $\eta^2 = 0.03$ | $\eta^2 = 0.03$ | $\eta^2 = 0.03$ |
| Aggressive Behavior | $\eta^2 = 0.03$ | $\eta^2 = 0.03$ | $\eta^2 = 0.03$ |
| Rulebreaking Behavior | $\eta^2 = 0.02$ | $\eta^2 = 0.02$ | $\eta^2 = 0.02$ |
| Intrusive Thoughts | $\eta^2 = 2.8 \times 10^{-3}$ | $\eta^2 = 3.9 \times 10^{-3}$ | $\eta^2 = 3.91 \times 10^{-3}$ |
| Depression (DSM) | $\eta^2 = 0.02$ | $\eta^2 = 0.02$ | $\eta^2 = 9.78 \times 10^{-3}$ |
| Anxiety Disorder (DSM) | $\eta^2 = 0.01$ | $\eta^2 = 9.46 \times 10^{-3}$ | $\eta^2 = 7.7 \times 10^{-4}$ |
| Somatic Problems (DSM) | $\eta^2 = 9.88 \times 10^{-3}$ | $\eta^2 = 0.01$ | $\eta^2 = 0.01$ |
| Avoidant (DSM) | $\eta^2 = 0.01$ | $\eta^2 = 0.01$ | $\eta^2 = 0.01$ |
| ADHD (DSM) | $\eta^2 = 0.03$ | $\eta^2 = 0.02$ | $\eta^2 = 0.03$ |
| Antisocial (DSM) | $\eta^2 = 0.03$ | $\eta^2 = 0.03$ | $\eta^2 = 0.03$ |
| Inattention (DSM) | $\eta^2 = 0.02$ | $\eta^2 = 0.03$ | $\eta^2 = 0.03$ |
| Hyperactivity/Impulsivity (DSM) | $\eta^2 = 0.02$ | $\eta^2 = 0.01$ | $\eta^2 = 0.01$ |
| Internalizing Problems | $\eta^2 = 0.03$ | $\eta^2 = 0.03$ | $\eta^2 = 0.03$ |
| Externalizing Problems | $\eta^2 = 0.04$ | $\eta^2 = 0.04$ | $\eta^2 = 0.04$ |
| Total Problems | $\eta^2 = 0.04$ | $\eta^2 = 0.05$ | $\eta^2 = 0.05$ |

**Table S7:** Abbreviation of Brain System Names.

| Abbreviated Name | Full Name |
| --- | --- |
| Aud | Auditory |
| CO | Cingulo-Opercular |
| CP | CentroParietal |
| DMN | Default Mode Network |
| DAN | Dorsal Attention Network |
| FP | Fronto-Parietal |
| None | None |
| RT | Retrosplenial-Temporal |
| SMh | Somatomotor (hand) |
| SMm | Somatomotor (mouth) |
| Sal | Salience |
| VAN | Ventral Attention Network |
| Vis | Visual |

**Figure S15:**
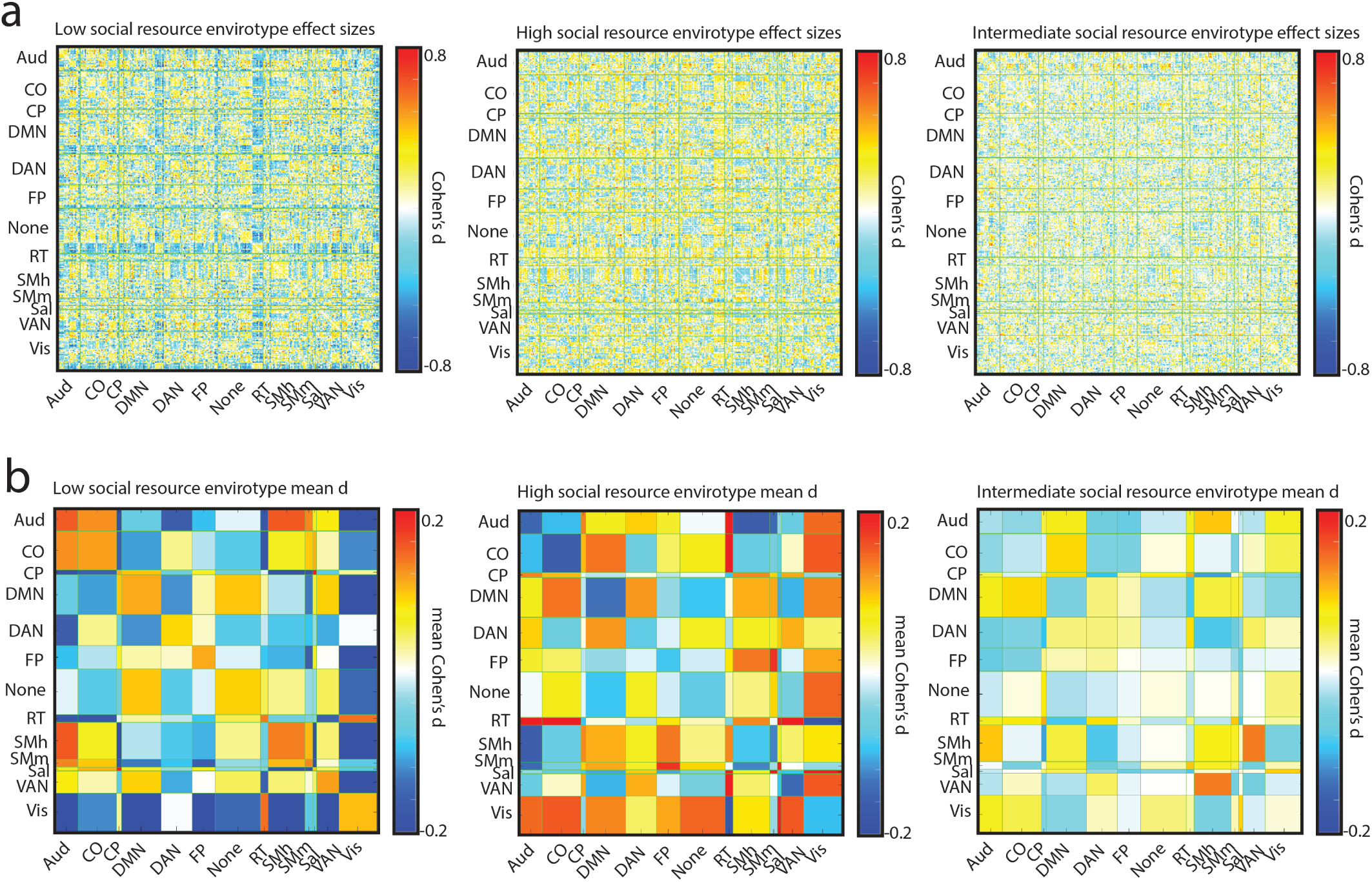
Effect sizes of differences in neural associations of externalizing problems across clusters. Row (a) shows Cohen’s *d* for all pairwise comparisons at the edge level. Row (b) depicts the mean Cohen’s *d* at the system level.

**Figure S16:**
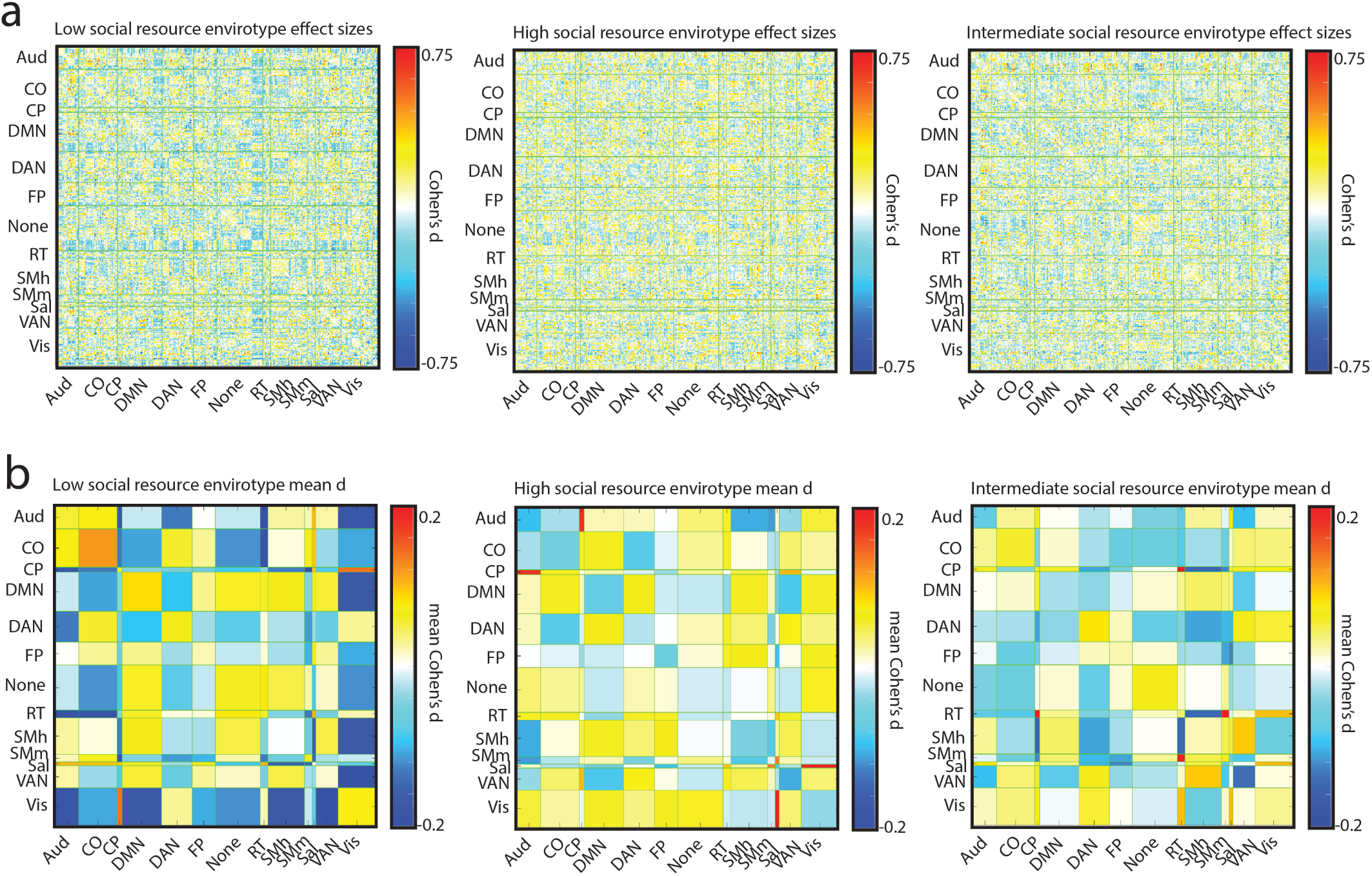
Effect sizes of differences in neural associations of internalizing problems across clusters. Row (a) shows Cohen’s *d* for all pairwise comparisons at the edge level. Row (b) depicts the mean Cohen’s *d* at the system level.

## Notes

### Competing Interest Statement

The authors have declared no competing interest.

## References

Amieva, H., Stoykova, R., Matharan, F., Helmer, C., Antonucci, T. C., and Dartigues, J.-F. (2010). What aspects of social network are protective for dementia? not the quantity but the quality of social interactions is protective up to 15 years later. Psychosomatic medicine, 72(9):905–911.

Antonucci, T. C., Ajrouch, K. J., and Birditt, K. S. (2014). The convoy model: Explaining social relations from a multidisciplinary perspective. The Gerontologist, 54(1):82–92.

Branchi, I. (2011). The double edged sword of neural plasticity: increasing serotonin levels leads to both greater vulnerability to depression and improved capacity to recover. Psychoneuroendocrinology, 36(3):339–351.

Bullmore, E. and Sporns, O. (2012). The economy of brain network organization. Nature Reviews Neuroscience, 13(5):336–349.

Casey, B. J., Cannonier, T., Conley, M. I., Cohen, A. O., Barch, D. M., Heitzeg, M. M., Soules, M. E., Teslovich, T., Dellarco, D. V., Garavan, H., et al. (2018). The adolescent brain cognitive development (abcd) study: imaging acquisition across 21 sites. Developmental cognitive neuroscience, 32:43–54.

Caspi, A. and Moffitt, T. E. (2006). Gene–environment interactions in psychiatry: joining forces with neuroscience. Nature Reviews Neuroscience, 7(7):583–590.

Coan, J. A., Beckes, L., Gonzalez, M. Z., Maresh, E. L., Brown, C. L., and Hasselmo, K. (2017). Relationship status and perceived support in the social regulation of neural responses to threat. Social Cognitive and Affective Neuroscience, 12(10):1574–1583.

Cocuzza, C. V., Chopra, S., Segal, A., Labache, L., Chin, R., Joss, K., and Holmes, A. J. (2026). Brain network dynamics reflect psychiatric illness status and transdiagnostic symptom profiles across health and disease. Nature Communications, 17(1):6678.

Cohen, S. (1988). Psychosocial models of the role of social support in the etiology of physical disease. Health psychology, 7(3):269.

Cohen, S. (2004). Social relationships and health. American psychologist, 59(8):676.

Cohen, S. and Wills, T. A. (1985). Stress, social support, and the buffering hypothesis. Psychological bulletin, 98(2):310.

Danese, A. and Widom, C. S. (2020). Objective and subjective experiences of child maltreatment and their relationships with psychopathology. Nature human behaviour, 4(8):811–818.

Dash, G. F., Karalunas, S. L., Kenyon, E. A., Carter, E. K., Mooney, M. A., Nigg, J. T., and Feldstein Ewing, S. W. (2023). Gene-by-environment interaction effects of social adversity on externalizing behavior in abcd youth. Behavior genetics, 53(3):219–231.

DeRosa, J., Friedman, N. P., Calhoun, V., and Banich, M. T. (2024). Neurodevelopmental subtypes of functional brain organization in the abcd study using a rigorous analytic framework. NeuroImage, 299:120827.

Drysdale, A. T., Grosenick, L., Downar, J., Dunlop, K., Mansouri, F., Meng, Y., Fetcho, R. N., Zebley, B., Oathes, D. J., Etkin, A., et al. (2017). Resting-state connectivity biomarkers define neurophysiological subtypes of depression. Nature medicine, 23(1):28–38.

Ellis, B. J. and Del Giudice, M. (2014). Beyond allostatic load: Rethinking the role of stress in regulating human development. Development and psychopathology, 26(1):1–20.

Ellwood-Lowe, M. E., Whitfield-Gabrieli, S., and Bunge, S. A. (2021). Brain network coupling associated with cognitive performance varies as a function of a child’s environment in the abcd study. Nature Communications, 12(1):1–14.

Feczko, E., Conan, G., Marek, S., Tervo-Clemmens, B., Cordova, M., Doyle, O., Earl, E., Perrone, A., Sturgeon, D., Klein, R., et al. (2021). Adolescent brain cognitive development (abcd) community mri collection and utilities. BioRxiv, pages 2021–07.

Fekson, V. K., Michaeli, T., Rosch, K. S., Schlaggar, B. L., and Horowitz-Kraus, T. (2023). Characterizing different cognitive and neurobiological profiles in a community sample of children using a non-parametric approach: An fmri study. Developmental Cognitive Neuroscience, 60:101198.

Foulkes, L. and Blakemore, S.-J. (2018). Studying individual differences in human adolescent brain development. Nature neuroscience, 21(3):315–323.

Garavan, H., Bartsch, H., Conway, K., Decastro, A., Goldstein, R., Heeringa, S., Jernigan, T., Potter, A., Thompson, W., and Zahs, D. (2018). Recruiting the abcd sample: Design considerations and procedures. Developmental cognitive neuroscience, 32:16–22.

Gee, D. G., Gabard-Durnam, L. J., Flannery, J., Goff, B., Humphreys, K. L., Telzer, E. H., Hare, T. A., Bookheimer, S. Y., and Tottenham, N. (2013). Early developmental emergence of human amygdala–prefrontal connectivity after maternal deprivation. Proceedings of the National Academy of Sciences, 110(39):15638–15643.

Glasser, M. F., Sotiropoulos, S. N., Wilson, J. A., Coalson, T. S., Fischl, B., Andersson, J. L., Xu, J., Jbabdi, S., Webster, M., Polimeni, J. R., et al. (2013). The minimal preprocessing pipelines for the human connectome project. Neuroimage, 80:105–124.

Gordon, E. M., Laumann, T. O., Adeyemo, B., Huckins, J. F., Kelley, W. M., and Petersen, S. E. (2016). Generation and evaluation of a cortical area parcellation from resting-state correlations. Cerebral cortex, 26(1):288–303.

Gordon, E. M., Laumann, T. O., Adeyemo, B., and Petersen, S. E. (2017). Individual variability of the system-level organization of the human brain. Cerebral cortex, 27(1):386–399.

Graham, S., Depp, C., Lee, E. E., Nebeker, C., Tu, X., Kim, H.-C., and Jeste, D. V. (2019). Artificial intelligence for mental health and mental illnesses: an overview. Current psychiatry reports, 21(11):116.

Heberle, A. E., Krill, S. C., Briggs-Gowan, M. J., and Carter, A. S. (2015). Predicting externalizing and internalizing behavior in kindergarten: Examining the buffering role of early social support. Journal of clinical child & adolescent psychology, 44(4):640–654.

Herrman, H., Patel, V., Kieling, C., Berk, M., Buchweitz, C., Cuijpers, P., Furukawa, T. A., Kessler, R. C., Kohrt, B. A., Maj, M., et al. (2022). Time for united action on depression: a lancet–world psychiatric association commission. The Lancet, 399(10328):957–1022.

Hicks, B. M., South, S. C., DiRago, A. C., Iacono, W. G., and McGue, M. (2009). Environmental adversity and increasing genetic risk for externalizing disorders. Archives of general psychiatry, 66(6):640–648.

Ho, T. C. and King, L. S. (2021). Mechanisms of neuroplasticity linking early adversity to depression: developmental considerations. Translational psychiatry, 11(1):517.

Holz, N. E., Zabihi, M., Kia, S. M., Monninger, M., Aggensteiner, P.-M., Siehl, S., Floris, D. L., Bokde, A. L., Desrivières, S., Flor, H., et al. (2023). A stable and replicable neural signature of lifespan adversity in the adult brain. Nature Neuroscience, 26(9):1603–1612.

Jirsaraie, R. J., Gatavins, M. M., Pines, A. R., Kandala, S., Bijsterbosch, J. D., Marek, S., Bogdan, R., Barch, D. M., and Sotiras, A. (2025). Mapping the neurodevelopmental predictors of psychopathology. Molecular Psychiatry, 30(2):478–488.

Krendl, A. C. and Perry, B. L. (2021). The impact of sheltering in place during the covid-19 pandemic on older adults’ social and mental well-being. The Journals of Gerontology: Series B, 76(2):e53–e58.

Lee, M. and Gonzalez, M. Z. (2025). Asymmetric access to social vs. economic resources during development calibrates socio-cognitive pathways to risk-taking in emerging adults. Cerebral Cortex, 35(7):bhaf169.

Lichenstein, S. D., Roos, C., Kohler, R., Kiluk, B., Carroll, K. M., Worhunsky, P. D., Witkiewitz, K., and Yip, S. W. (2022). Identification and validation of distinct latent neurodevelopmental profiles in the adolescent brain and cognitive development study. Biological Psychiatry: Cognitive Neuroscience and Neuroimaging, 7(4):352–361.

Liu, P., Song, D., Guo, Y., and Zhang, H. (2025). Brain functional connectivity mediates the association between adverse childhood experiences and conduct problems. Biological Psychiatry: Cognitive Neuroscience and Neuroimaging.

Lynch, C. J., Elbau, I. G., Ng, T., Ayaz, A., Zhu, S., Wolk, D., Manfredi, N., Johnson, M., Chang, M., Chou, J., et al. (2024). Frontostriatal salience network expansion in individuals in depression. Nature, 633(8030):624–633.

McLaughlin, K. A., Sheridan, M. A., and Lambert, H. K. (2014). Childhood adversity and neural development: deprivation and threat as distinct dimensions of early experience. Neuroscience & Biobehavioral Reviews, 47:578–591.

Merritt, H., Faskowitz, J., Gonzalez, M. Z., and Betzel, R. F. (2024). Stability and variation of brain-behavior correlation patterns across measures of social support. Imaging Neuroscience, 2:1–18.

Merritt, H., Koch, M. K., Jo, Y., Chumin, E. J., and Betzel, R. F. (2026a). Social ‘envirotyping’the abcd study contextualizes dissociable brain organization and diverging outcomes. Social Cognitive Affective Neuroscience.

Merritt, H., Rakesh, D., and Betzel, R. (2026b). Connection context: The variable neural architecture of social relationships. Neuroscience and Biobehavioral Reviews, 191.

Michael, C., Larsen, B., Satterthwaite, T. D., and Hyde, L. W. (2025). Mapping interactions between adversity and neuroplasticity across development. Trends in Cognitive Sciences.

Morawetz, C., Berboth, S., and Bode, S. (2021). With a little help from my friends: The effect of social proximity on emotion regulation-related brain activity. Neuroimage, 230:117817.

Mwilambwe-Tshilobo, L., Ge, T., Chong, M., Ferguson, M. A., Misic, B., Burrow, A. L., Leahy, R. M., and Spreng, R. N. (2019). Loneliness and meaning in life are reflected in the intrinsic network architecture of the brain. Social cognitive and aflective neuroscience, 14(4):423–433.

Mwilambwe-Tshilobo, L., Setton, R., Bzdok, D., Turner, G. R., and Spreng, R. N. (2022). Age differences in functional brain networks associated with loneliness and empathy. Network Neuroscience, pages 1–60.

Prompiengchai, S. and Dunlop, K. (2024). Breakthroughs and challenges for generating brain network-based biomarkers of treatment response in depression. Neuropsychopharmacology, pages 1–16.

Rajaleid, K., Nummi, T., Westerlund, H., Virtanen, P., Gustafsson, P. E., and Hammarström, A. (2016). Social adversities in adolescence predict unfavourable trajectories of internalized mental health symptoms until middle age: results from the northern swedish cohort. The European Journal of Public Health, 26(1):23–29.

Rakesh, D., Allen, N. B., and Whittle, S. (2021a). Longitudinal changes in within-salience network functional connectivity mediate the relationship between childhood abuse and neglect, and mental health during adolescence. Psychological medicine, pages 1–13.

Rakesh, D., Kelly, C., Vijayakumar, N., Zalesky, A., Allen, N. B., and Whittle, S. (2021b). Unraveling the consequences of childhood maltreatment: deviations from typical functional neurodevelopment mediate the relationship between maltreatment history and depressive symptoms. Biological psychiatry: cognitive neuroscience and neuroimaging, 6(3):329–342.

Rakesh, D., Sadikova, E., and McLaughlin, K. A. (2025a). Associations among socioeconomic disadvantage, longitudinal changes in within-network connectivity, and academic outcomes in the abcd study. Developmental Cognitive Neuroscience, page 101587.

Rakesh, D., Seguin, C., Zalesky, A., Cropley, V., and Whittle, S. (2021c). Associations between neighborhood disadvantage, resting-state functional connectivity, and behavior in the adolescent brain cognitive development study: the moderating role of positive family and school environments. Biological Psychiatry: Cognitive Neuroscience and Neuroimaging, 6(9):877–886.

Rakesh, D., Tsomokos, D. I., Vargas, T., Pickett, K. E., and Patel, V. (2025b). Macroeconomic income inequality, brain structure and function, and mental health. Nature Mental Health, 3(11):1318–1330.

Rakesh, D. and Whittle, S. (2021). Socioeconomic status and the developing brain–a systematic review of neuroimaging findings in youth. Neuroscience & Biobehavioral Reviews, 130:379–407.

Rakesh, D., Whittle, S., Sheridan, M. A., and McLaughlin, K. A. (2023). Childhood socioeconomic status and the pace of structural neurodevelopment: accelerated, delayed, or simply different? Trends in Cognitive Sciences.

Rakesh, D., Zalesky, A., and Whittle, S. (2021d). Similar but distinct–effects of different socioeconomic indicators on resting state functional connectivity: Findings from the adolescent brain cognitive development (abcd) study®. Developmental cognitive neuroscience, 51:101005.

Ramphal, B., DeSerisy, M., Pagliaccio, D., Raffanello, E., Rauh, V., Tau, G., Posner, J., Marsh, R., and Margolis, A. E. (2020). Associations between amygdala-prefrontal functional connectivity and age depend on neighborhood socioeconomic status. Cerebral Cortex Communications, 1(1):tgaa033.

Ruiz, W. D. and Yabut, H. J. (2024). Autonomy and identity: the role of two developmental tasks on adolescent’s wellbeing. Frontiers in Psychology, 15:1309690.

Segal, A., Tiego, J., Parkes, L., Holmes, A. J., Marquand, A. F., and Fornito, A. (2025). Embracing variability in the search for biological mechanisms of psychiatric illness. Trends in Cognitive Sciences, 29(1):85–99.

Sheridan, M. A. and McLaughlin, K. A. (2014). Dimensions of early experience and neural development: deprivation and threat. Trends in cognitive sciences, 18(11):580–585.

Spreng, R. N., Dimas, E., Mwilambwe-Tshilobo, L., Dagher, A., Koellinger, P., Nave, G., Ong, A., Kernbach, J. M., Wiecki, T. V., Ge, T., et al. (2020). The default network of the human brain is associated with perceived social isolation. Nature communications, 11(1):1–11.

Sydnor, V. J., Larsen, B., Seidlitz, J., Adebimpe, A., Alexander-Bloch, A. F., Bassett, D. S., Bertolero, M. A., Cieslak, M., Covitz, S., Fan, Y., et al. (2023). Intrinsic activity development unfolds along a sensorimotor–association cortical axis in youth. Nature Neuroscience, pages 1–12.

Tooley, U. A., Bassett, D. S., and Mackey, A. P. (2021). Environmental influences on the pace of brain development. Nature Reviews Neuroscience, 22(6):372–384.

Tottenham, N. (2014). The importance of early experiences for neuro-affective development. The neurobiology of childhood, pages 109–129.

Vaidya, N., Marquand, A. F., Nees, F., Siehl, S., and Schumann, G. (2024). The impact of psychosocial adversity on brain and behaviour: an overview of existing knowledge and directions for future research. Molecular psychiatry, 29(10):3245–3267.

Vanes, L., Rakesh, D., Banaschewski, T., Bodke, A. L., Desrivières, S., Flor, H., Garavan, H., Gowland, P., Grigis, A., Heinz, A., et al. (2025). Longitudinal associations of structural and functional brain connectivity with dimensions of psychopathology in adolescence. Biological Psychiatry: Cognitive Neuroscience and Neuroimaging.

Voepel-Lewis, T., Stoddard, S. A., Ploutz-Snyder, R. J., Chen, B., and Boyd, C. J. (2025). Effect of comorbid psychologic and somatic symptom trajectories on early onset substance use among us youth in the abcd study. Addictive behaviors, 160:108181.

Wang, Z., Zhou, X., Gui, Y., Liu, M., and Lu, H. (2023). Multiple measurement analysis of resting-state fmri for adhd classification in adolescent brain from the abcd study. Translational Psychiatry, 13(1):45.

Webb, E. K., Cardenas-Iniguez, C., and Douglas, R. (2022). Radically reframing studies on neurobiology and socioeconomic circumstances: A call for social justice-oriented neuroscience. Frontiers in Integrative Neuroscience, 16.

Whittle, S., Zhang, L., and Rakesh, D. (2025). Environmental and neurodevelopmental contributors to youth mental illness. Neuropsychopharmacology, 50(1):201–210.

Wong, Y. J., Rew, L., and Slaikeu, K. D. (2006). A systematic review of recent research on adolescent religiosity/spirituality and mental health. Issues in mental health nursing, 27(2):161–183.

Xiao, M., Chen, X., Yi, H., Luo, Y., Yan, Q., Feng, T., He, Q., Lei, X., Qiu, J., and Chen, H. (2021). Stronger functional network connectivity and social support buffer against negative affect during the covid-19 outbreak and after the pandemic peak. Neurobiology of stress, 15:100418.

Xie, C., Xiang, S., Shen, C., Peng, X., Kang, J., Li, Y., Cheng, W., He, S., Bobou, M., Broulidakis, M. J., et al. (2023). A shared neural basis underlying psychiatric comorbidity. Nature medicine, 29(5):1232–1242.

